# MSqRob takes the missing hurdle: uniting intensity- and count-based proteomics

**DOI:** 10.1101/782466

**Authors:** Ludger J.E. Goeminne, Adriaan Sticker, Lennart Martens, Kris Gevaert, Lieven Clement

## Abstract

Missing values are a major issue in quantitative data-dependent mass spectrometry-based proteomics. We therefore present an innovative solution to this key issue by introducing a hurdle model, which is a mixture between a binomial peptide count and a peptide intensity-based model component. It enables dramatically enhanced quantification of proteins with many missing values without having to resort to harmful assumptions for missingness. We demonstrate the superior performance of our method by comparing it with state-of-the-art methods in the field.

---

Label-free data-dependent quantitative mass spectrometry (MS) is the preferred method for deep and high-throughput identification and quantification of thousands of proteins in a single analysis^1^. However, this approach suffers from many missing values, which strongly reduce the amount of quantifiable proteins^2, 3^. There are three common causes for this missingness: (1) true absence of signal, or signal below detection limit in the MS1 spectrum; (2) lack of fragmentation and hence missed identification of the MS1 peak; and (3) failed identification of the acquired fragmentation spectrum. As a result, missingness is more likely to occur for low-abundant proteins and/or poorly ionizing peptides. However, missingness may also extend to mid- and even high-range intensities, e.g. when co-eluting peptides suppress an MS1 signal or when poor quality of an MS2 spectrum interferes with correct identification. Missingness due to lack of fragmentation can be mitigated by “matching between runs”, where unidentified MS1 peaks in one run are aligned to identified peaks in another run in narrow retention time and mass-over-charge (*m*/*z*) windows^4^. Nevertheless, missing values remain widespread; a survey of 73 recent public proteomics data sets from the PRIDE database demonstrates an average of 44% missing values (Fig. 1 a, Supplementary Table 1).

**Figure 1.**
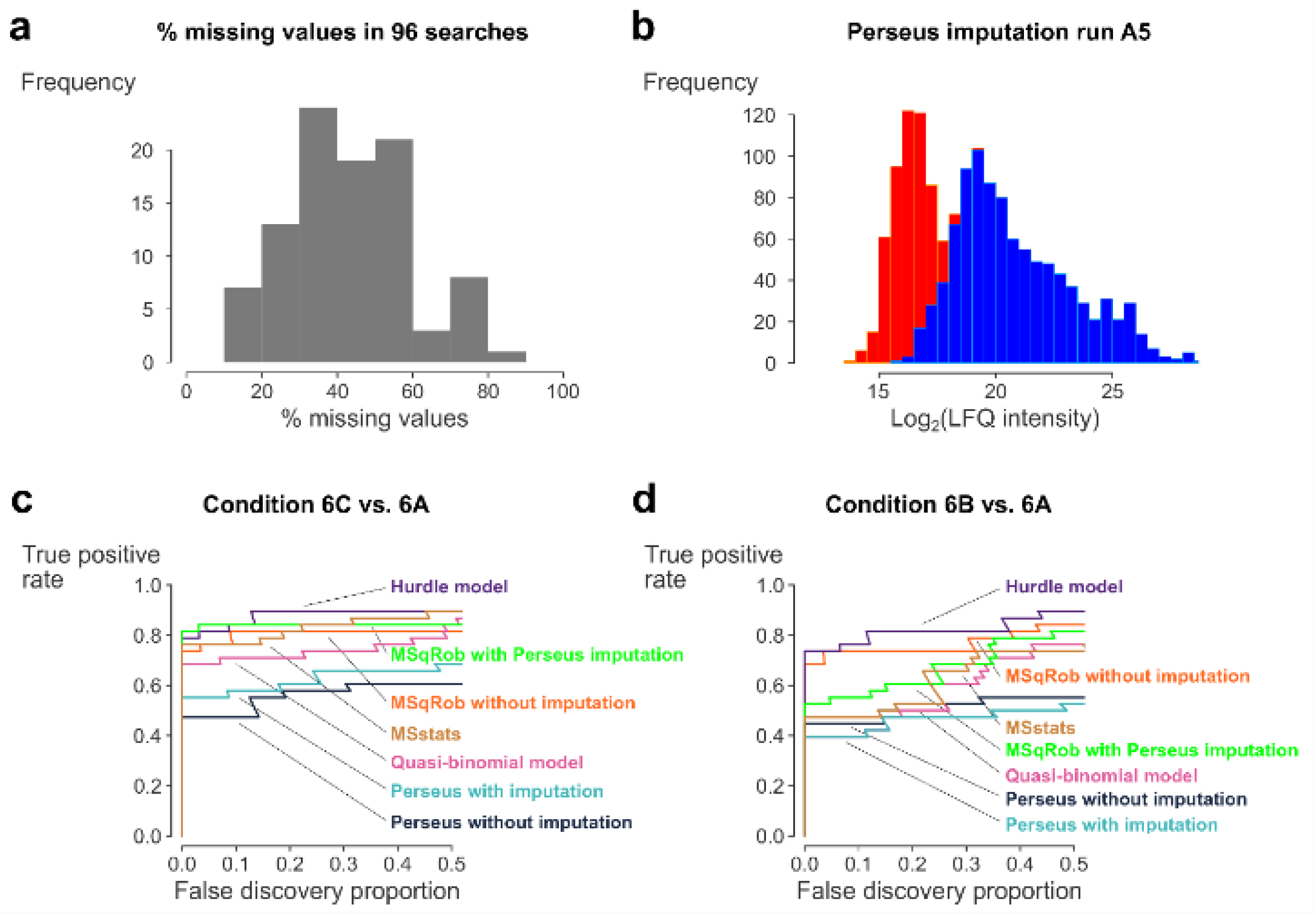
The impact of missing values and the superior performance of the hurdle model. (a) Missing values are highly prevalent in recent proteomics data. The histogram shows the distribution of the percentage of missing values in 96 peptides.txt files from 73 PRIDE projects with a PRIDE publication date in 2017 that applied shotgun proteomics to full or partial proteomes. Missingness ranges from 16 to 82%, with an average of 44% missing values. (b) Perseus imputation fails when too many missing values are present: the frequency distribution of the log_2_-transformed LFQ intensities in the CPTAC dataset becomes bimodal after Perseus imputation due to the many imputed values, as exemplified by run A5. Observed values (978 LFQ intensities) are colored in blue, imputed values (484 LFQ intensities) are colored in red. Similar results for the other runs can be seen in Supplementary Fig. 1. (c, d) In the CPTAC dataset (38 UPS1 and 1,343 yeast proteins), the hurdle model outperforms other methods. The fraction of true positive UPS1 proteins flagged as DA in conditions C vs. A and B vs. A is plotted as a function of the fraction of false positive yeast proteins in the total number of DA proteins.

A common solution for missingness is to impute the missing values. But, this comes at the expense of additional assumptions, e.g. k-nearest neighbors (kNN) assumes intensity-independent missingness, and the popular software packages Perseus^5^ (“Perseus imputation”, PI) and MSstats^6^ assume missingness by low intensity. In reality, however, missing values originate from a mix of intensity-dependent and -independent mechanisms, which can moreover be strongly data set-specific^3, 7^. As a result, state-of-the-art imputation profoundly changes the distribution of protein-level intensities as it typically produces a second mode due to many imputed values, (shown for MaxQuant/Perseus in Fig. 1 b and Supplementary Fig. 1 – 2; and for kNN in Supplementary Fig. 3 – 4). This can have a deep impact on the downstream differential analysis. We demonstrate these effects on the widely studied CPTAC dataset^8^, where more than 47% of peptide data are missing. Because the spike-in concentrations in this dataset cover a wide range, the perils of imputation can be clearly shown. Indeed, while our MSqRob^9^ quantification tool shows excellent performance when combined with PI for comparison C (high spike-in concentration: 2.2 fmol/µL) vs. A (lowest spike in concentration: 0.25 fmol/µL) (Fig. 1 c), MSqRob with PI performs poorly for comparisons B (intermediate amount of spike-in: 0.74 fmol/µL) vs. A (Fig. 1 d) and C vs. B, where the differences in spike-in concentration are smaller (Supplementary Fig. 5, 6), thus rapidly accumulating false positives. MSqRob therefore omits imputation by default to avoid a severe backlash in performance^10^. However, without imputation, intensity-based methods cannot cope with complete missingness in one condition as it is impossible to calculate a fold change (FC) relative to a missing value. Thus, potentially interesting cases such as strong protein synthesis/stabilization or protein degradation remain undetected.

Conversely, peptide counting approaches naturally handle missing peptides in a run by a zero count. Yet relative quantification by peptide counting generally performs poorly (pink line in Fig. 1 c, d). This because counting disregards the inherent abundance-intensity relationship (within a given dynamic range) for each peptide^11, 12^. However, differentially abundant (DA) proteins for which no FC can be estimated due to missingness do differ in peptide counts (Supplementary Fig. 7) and proteins with a higher concentration have on average both higher peptide ion intensities and higher peptide counts, as it is more likely that even their poorly ionizing peptides are detected.

Combining intensity-based methods with counting-based methods to exploit this complementary information therefore seems promising. Webb-Robertson *et al.* combine intensity- and count-based statistics to filter out peptides prior to differential analysis^13^. ProPCA uses principal components to summarize intensities and spectral counts into one value^14^ and IDPQuantify uses Fisher’s method to combine the p-values from a two-sample t-test with those of a quasi-Poisson regression on spectral counts^15^. However, ProPCA’s combined metric lacks an intuitive interpretation, and does not include any downstream statistical analysis. Like Perseus, IDPQuantify only handles simple pairwise comparisons without any possibility to correct for confounding effects. Moreover, none of these methods is able to pinpoint whether the statistical significance is driven by the count component, the intensity component, or both, thus making it impossible to interpret the results in terms of FCs.

Here, we introduce a hurdle model that unites the advantages of MSqRob with the complementary information present in peptide counts, that avoids unrealistic imputation assumptions, and that provides interpretable results. This hurdle model takes peptide-level information as input and consists of a mixture model of two components: (1) a binary component that distinguishes between log_2_-transformed peptide intensities that are either missing or observed; and (2) an MSqRob-based component to model the magnitude of log_2_ peptide intensities passing the detection hurdle. Inference on the parameters of both model components has an intuitive interpretation: the binary component can be used to assess *differential detection (DD)* and returns *log odds ratios* (log ORs), while the MSqRob component allows to test for *differential abundance (DA)* of a protein and returns *log*_*2*_ *fold changes* (log_2_ FCs). When peptides are completely missing in one condition, the hurdle model reduces to a binomial model. However, when peptides are present in both conditions, the hurdle approach allows to combine information in the OR and FC test statistics, and can be used to infer on DD, DA, or both in a post-hoc analysis. Note, that the peptide counts can be overdispersed, which is why the variance component is estimated via quasi-likelihood and the model is termed a quasi-binomial model.

## EXPERIMENTAL SECTION

In this section, we first demonstrate how we analyzed the frequency of missing values in recent PRIDE projects. Then, we discuss the nature and the origin of the CPTAC and the HEART datasets, followed by an overview of how these datasets were preprocessed and imputed with different strategies. We then introduce the Perseus, MSstats, MSqRob, the quasi-binomial model and the hurdle model as methods for statistical inference. Finally, we refer to our GitHub page which enables to reproduce all the analyses in this publication.

### Missing values in recent PRIDE projects

For our analysis of the frequency of missing values of recent datasets in PRIDE, we downloaded 96 peptides.txt files corresponding to 73 PRIDE projects that adhere to the following conditions:

- label-free shotgun proteomics,
- full proteomes or enriched subsets of the proteome (i.e. no projects where there was an enrichment step for a chemical modification and no projects investigating protein-protein interactions),
- published in PRIDE in 2017 and
- searched with MaxQuant and peptides.txt file available from PRIDE.

An overview of the projects and peptides.txt files with their number of missing values, number of observed values and percentage of missing values is given in Supplementary Table 1.

### Data availability

We made use of two example datasets.

1. In the benchmark CPTAC study 6, which can be downloaded from https://cptac-data-portal.georgetown.edu/cptac/dataPublic/list?currentPath=%2FPhase_I_Data%2FStudy6&nonav=true, the human UPS1 standard is spiked in 5 different concentrations in a yeast proteome background^8^. This allows to make comparisons for which the ground truth is known: when comparing two different spike-in conditions, only the UPS1 proteins are truly differentially abundant (DA), while the yeast proteins are not.

The three lowest spike-in conditions (A, B and C) from LTQ-Orbitrap at site 86, LTQ-Orbitrap O at site 65 and LTQ-Orbitrap W at site 56 were searched with MaxQuant 1.6.1.0 against a database containing 6,718 reviewed *Saccharomyces cerevisiae* (strain ATCC 204508/S288c) proteins downloaded from UniProt on September 14, 2017, supplemented with the 48 human UPS1 protein sequences provided by Sigma Aldrich. Carbamidomethylcysteine was set as a fixed modification and methionine oxidation, protein N-terminal acetylation and N-terminal glutamine to pyroglutamate conversion were set as variable modifications. Detailed search settings are described in Supplementary Material.

We only performed pairwise comparisons between the 3 lowest spike-in concentrations because of the huge ionization competition effects in the higher spike-in concentrations, as described earlier^9^.

2. The HEART dataset originates from a large-scale proteomics study of the human heart, where 16 different regions and 3 different cell types from three healthy human adult hearts were studied^16^.

For our analysis, we made use of the peptides.txt and proteinGroups.txt files made available by Doll *et al.* in ProteomeXchange via the PRIDE Archive repository under the identifier PXD006675. Here, we limited ourselves to the data from 6 regions: the atrial and ventricular septa (SepA and SepV), the left and right atrium (LA and RA) and the left and right ventricle (LV and RV). We compare the atrial regions (LA, RA and SepA) to the ventricular regions (LV, RV and SepV), as well as LA vs. RA and LV vs. RV.

### Preprocessing for MSqRob and the quasi-binomial model

Peptide intensities obtained from MaxQuant’s peptides.txt file were log_2_-transformed and quantile normalized. Next, potential contaminants, reverse sequences and proteins that were only identified by peptides carrying a modification were removed from the data. Finally, proteins identified by only a single peptide were also removed.

### Imputation methods

Missing values were either not imputed or imputed with kNN or Perseus imputation.

#### No imputation

For MSqRob without imputation, we additionally removed peptides that are only identified in a single run. Indeed, MSqRob directly models peptide intensities and the peptide-specific effect for peptides with one identification cannot be estimated because no replicates are available. This additional filtering step is not required for MSqRob with imputation, because missing values in the peptide intensity matrix are imputed first.

#### kNN imputation

k-nearest neighbors (kNN) imputation calculates a Euclidean distance metric on all peptide intensities to find the *k* most similar peptides and imputes the missing value with the average of the corresponding values from the *k* neighbors^17^. We used the default value of *k* = 10 neighbors.

#### Perseus imputation

We call “Perseus imputation (PI)” the standard imputation approach from the popular proteomics computational platform Perseus^5^. The imputation is achieved by using the “Replace missing values from normal distribution” function in Perseus 1.0.6.7. We applied PI on peptide intensities for MSqRob with PI and on LFQ intensities for Perseus with PI.

PI first constructs a rescaled normal distribution with: (1) a downshifted mean equal to the average of all the observed data minus *d* times the standard deviation of the observed data and (2) a standard deviation equal to *w* times the standard deviation of the observed data. Missing values are then imputed with random draws from this rescaled distribution. We adopt Perseus’ default values for *w* (0.3) and *d* (1.8).

### Statistical inference

In this subsection, we discuss differential protein analysis with Perseus and MSstats, followed by MSqRob, the quasi-binomial model and our hurdle model. All methods mentioned below model the data protein by protein. To improve readability, we will suppress the protein indicator in the remainder of this section.

#### Perseus

For the CPTAC dataset, LFQ intensities were imported into Perseus 1.6.0.7. Potential contaminants, reversed sequences and proteins only identified by peptides carrying modification sites were removed from the data. Next, empty columns were removed and via “Categorical annotation rows”, runs were sorted according to their spike-in conditions (A, B or C). Data were either imputed with PI (“Perseus with imputation”) or not (“Perseus without imputation”). Finally, we performed two-sample t-tests for each of the three comparisons.

For the atrial vs. ventricular comparison in the HEART dataset, Doll *et al.* provided the results of their Perseus with imputation approach in their Supplementary Data 5 file, available from https://static-content.springer.com/esm/art%3A10.1038%2Fs41467-017-01747-2/MediaObjects/41467_2017_1747_MOESM7_ESM.xlsx.

#### MSqRob

MSqRob has been described in detail in Goeminne *et al.* (2016)^9^. Briefly, for each protein, the log_2_-transformed peptide intensities *y*_*pr*_ for peptide *p* = 1, …, *P* in run *r* = 1, …, *R* are modeled as follows:

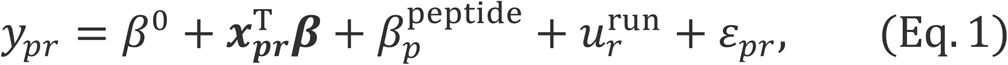

where *β*^0^ is the intercept, 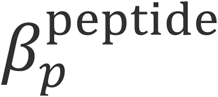 is the fixed effect of the individual peptide sequence *p*, 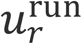 is a random run effect 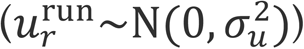 that accounts for the correlation of peptide intensities *y*_*jr*_ and *y*_*kr*_ from the same protein within the same run *r*, 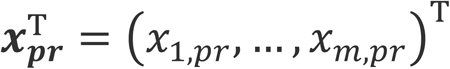 is a vector with the covariate pattern of the *m* remaining predictors, ***β*** = [*β*_1_, …, *β*_*m*_]^T^ is a vector of parameters modeling the effect of each predictor on the peptide intensity conditionally on the remaining covariates, and *ε*_*pr*_ is the error term that is assumed to be normally distributed (*ε*_*pr*_∼N(0, *σ*^2^)).

For the CPTAC and HEART datasets, the MSqRob model can be written as follows:

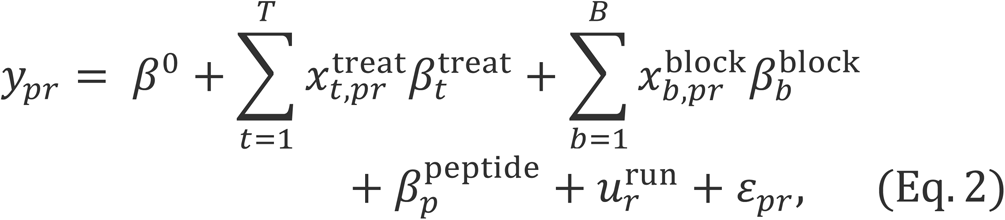

where *β*^0^ is the intercept, 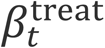 is the effect of interest (*t* = 1 to 3 corresponding to spike-in conditions A, B, C for CPTAC, *t* = 1 to 6 for the six cardiac regions in the HEART dataset), 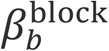 is a blocking factor (lab *b* = 1 to 3 for CPTAC, patient *b* = 1 to 3 for HEART), 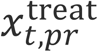 and 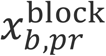 are dummy variables which are equal to 1 if run *r* corresponds to treatment *t* or block *b*, respectively, and 0 otherwise. 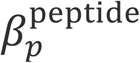 is the effect of the individual peptide sequence *p*, 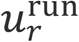 is a random effect that accounts for the fact that peptides within each run *r* are correlated. Due to the parameterization of the model, the following restrictions apply: 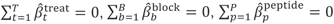 and 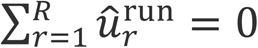. The effect sizes 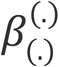 (except for the intercept) are estimated using penalized regression. Distinct ridge penalties are used for the treatment, block and peptide parameters, respectively and the ridge penalties are tuned by exploiting the link between ridge regression and mixed models (see e.g. Ruppert *et al.* (2003), chapter 4^18^). Outliers are accounted for using M-estimation with Huber weights. Protein-wise degrees of freedom are now calculated in a less liberal way as: *R* − (1 + (*T* − 1) + (*B* − 1)). Variances are stabilized by borrowing information over proteins using limma’s empirical Bayes approach, which results in a moderated t-test. Scripts to run MSqRob are provided on our GitHub page (see below under “Code availability”).

#### MSstats

MSstats has been described by Choi *et al.* (2014)^6^. For our analysis, we used the default settings of MSstats version 3.12.2. During preprocessing, feature intensities are log_2_-transformed and normalized by equalizing the run medians. Then, missing values are imputed with the default MSstats settings. Features are summarized to the protein level with Tukey’s median polish method. Treatment effects are specified as “Condition” and blocking factors as “BioReplicate” in the Msstats workflow.

#### Quasi-binomial model

When assuming that the number of observed peptides 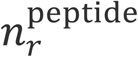 in each run *r* (after preprocessing) for a protein are binomially distributed

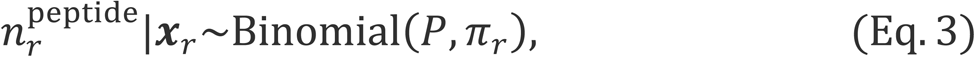

with *P* the total number of unique peptides observed over all runs *r* = 1, …, *R* for this protein, and *π*_*r*_ the probability to identify a peptide for this protein; the peptide counts can be modeled using logistic regression. However, they are often overdispersed with respect to the binomial distribution. We therefore adopt quasi-binomial regression (McCullagh and Nelder (1989), section 4.5^19^) where we model the first two moments (mean 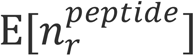 and variance 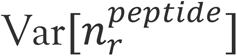) as follows:

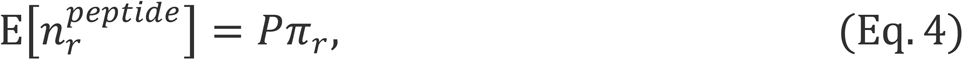

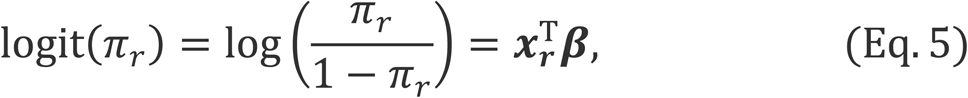

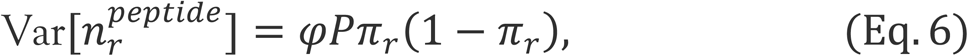

with *φ* a dispersion parameter accommodating for a more flexible variance function than that of the binomial regression model. Similar to the original edgeR count model for gene expression, information is borrowed over the dispersion parameters of all models with the empirical Bayes procedure of the limma R package^20^. To avoid overly liberal results from inaccurate variance estimations, underdispersed *φ* estimates are set to 1. Note that the quasi-binomial approach models the log odds, i.e. the logarithm of the probability that a peptide is detected divided by the probability that a peptide is not detected, i.e. the odds on detection. Hence, the model will return log ORs when contrasting different treatments to each other, which can be used to infer on differential peptide detection for a specific protein.

#### Hurdle model

The normalized intensities for each peptide *p* in each run *r* are typically assumed to follow a log-normal distribution. Upon log_2_-transformation, missing (zero) intensities are set at −∞ and cannot be modeled with intensity-based methods such as Perseus, MSstats and MSqRob. Missing values are therefore either omitted or imputed.

Here, we consider a hurdle model that consists of two parts: a binary component *z*_*pr*_ that distinguishes between peptide intensities in run *r* that are missing (*z*_*pr*_ = 0) or observed (*z*_*pr*_ = 1) with detection probability *π*_*r*_; and a normal component *y*_*pr*_ with mean *μ*_*pr*_ and variance *σ*^2^ to model log_2_ peptide intensities passing the detection hurdle. Note, that the detection probability *π*_*r*_ and the mean *μ*_*p r*_ can be further parameterized using peptide specific effects, a random run effect 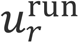, and additional covariates ***x***_*pr*_. More formally, the hurdle model for log_2_-transformed intensities *y*_*pr*_ for peptide *p* = 1, …, *P* in run *r* = 1, …, *R* can be specified as follows:

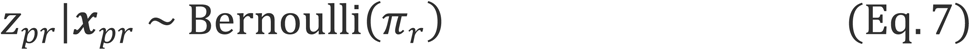

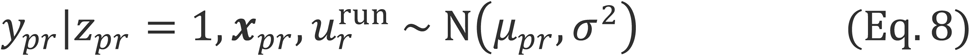

The log-likelihood for ***π*** = [*π*_1_, …, *π*_*R*_]^T^, ***μ*** = [*μ*_11_, …, *μ*_*PR*_]^T^ and *σ*^2^ given ***y*** = [*y*_11_, …, *y*_*PR*_]^T^, ***z*** = [*z*_11_, …, *z*_*PR*_]^T^, ***u***^*run*^ = [*u*_1_, …, *u*_*R*_]^T^, and ***X*** the matrix with rows 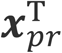 can then be written as:

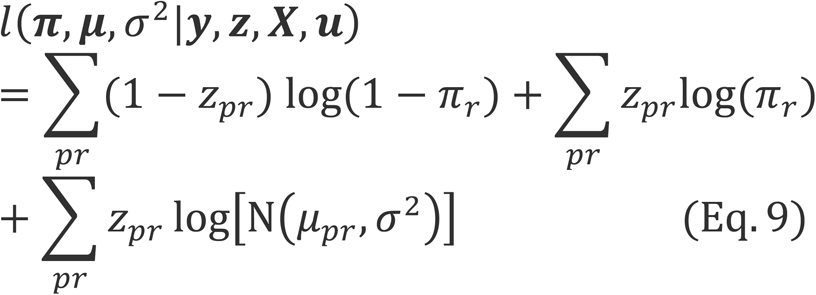

Note, that the log-likelihood implies an estimation orthogonality between *π*_*r*_ and *μ*_*pr*_ and that the first two terms in the equation are equivalent to the log-likelihood of a Bernoulli process. Further, we omit a peptide-specific effect for the parameterization of *π*_*r*_ because this leads to complete separation for too many proteins. The detection probability is thus considered constant for all peptides of a particular protein in run *r*. When summing over the peptides *p* = 1, …, *P*, peptide counts 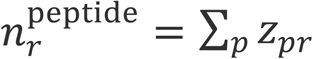 are obtained and it is computationally more efficient to estimate *π*_*r*_ using a binomial model for the peptide counts. To account for overdispersion in the counts, we will again estimate the detection probability *π*_*r*_ using quasi-binomial regression with Model (Eq. 4 − 6). The last term in the log-likelihood corresponds to a normal log-likelihood and we propose to estimate *μ*_*pr*_ and *σ*^2^ using the MSqRob Model (Eq. 2).

Both model components model their means using the same covariate pattern ***x***_*pr*_ and they will allow us to assess the same contrast of interest to infer on the log OR on detection and a log_2_ FC between conditions, respectively. Assuming independence between both statistics under the null hypothesis (Supplementary Table 5, Supplementary Figure 14), the hurdle model allows to assess the omnibus null hypothesis of no differential detection and no differential expression by combining the p-values of MSqRob without imputation and of the quasi-binomial model component. To this end, we first transform the p-values to z-values and combine them in a chi-square statistic:

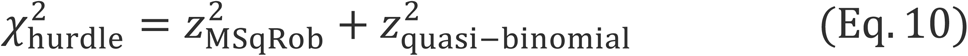

If an MSqRob log_2_ FC can be estimated the chi-square statistic follows a chi-square distribution with 2 degrees of freedom under the omnibus null hypothesis, otherwise the corresponding p-value is equivalent to that of the quasi-binomial model.

#### Two-stage inference

We used stageR version 1.3.29 to implement a two-stage inference procedure^21^. It is a two-stage testing paradigm that leverages power of aggregating multiple tests per protein (here a test for differential detection, DD and differential abundance, DA) in the screening stage. Upon rejection of the omnibus null hypothesis, stageR performs post-hoc tests to assess on DD and DA. We adopt the Benjamini-Hochberg False Discovery Rate (FDR) procedure on the aggregated tests in the first stage. In the post-hoc analysis, we use the modified Holm procedure implemented in stageR to control the Family Wise Error Rate for the DD and DA tests within a protein at the FDR-adjusted significance level of the first stage.

#### Multiple testing correction

All p-values for MSstats, MSqRob, the quasi-binomial and the hurdle model were corrected for multiple testing using the Benjamini-Hochberg FDR correction. Perseus uses a permutation-based FDR based on 250 iterations.

## Code availability

All scripts to reproduce the results in this contribution and in supplementary material are available at https://github.com/statOmics/MSqRobHurdlePaper.

## RESULTS AND DISCUSSION

We use the CPTAC study 6 dataset^8^ to compare our hurdle approach to the quasi-binomial model alone, to MSqRob alone, to Perseus (with and without imputation), and to MSstats. Fig. 1 c, d demonstrates the superior performance of the hurdle model: it consistently outperforms the other algorithms. Moreover, the hurdle model also always outperforms MSqRob with imputation under a low abundance assumption such as PI (Supplementary Fig. 5, 6). In the lowest spike-in condition, A, missing values in the UPS1 proteins are mainly caused by low abundance, allowing the count-component of the hurdle model to add to the strength of the intensity-component. In comparison C vs. B, the UPS1 spike-in concentrations are higher, reducing the difference in peptide counts between both conditions, and mainly driving significance of the hurdle model by the difference in average intensities. In this case, the hurdle model performs on par with MSqRob without imputation, as is expected, while PI approaches still perform poorly (Supplementary Fig. 5, 6).

Next, we assess the human heart dataset of Doll *et al.* (2017)^16^. For the 7,822 gene identifiers in common after preprocessing, we assessed the overlap between the 1,500 most significantly regulated identifiers in the Doll *et al.*, hurdle, and MSqRob comparisons between the atrial to the ventricular proteome (Fig. 2 a, Supplementary Table 2). The 671 identifiers shared between all methods correspond to proteins with a strong DA and many identified peptides (Supplementary Fig. 8). The 158 identifiers shared between Perseus and hurdle mostly have strong DD and only few peptides in one of the heart chambers (Supplementary Fig. 9). The 164 identifiers unique to hurdle differ strongly in peptide counts, but the log_2_ FCs are close to one. (Supplementary Fig. 10). The 596 identifiers unique to Perseus often show very few peptide identifications or very small FCs, making these results more questionable (Supplementary Fig. 11). Indeed, the imputation strategy again has a profound impact on the results: there is a 22% non-overlap between Perseus with and without the PI strategy. Moreover, because of the stochastic nature of the PI imputation, on average 4.5% of the first 1,500 proteins declared DA did not overlap between two repeated PI analyses. The 75 identifiers shared between MSqRob and Perseus, finally, mainly have many peptides, but a relatively small FC (Supplementary Fig. 12). These are not included in the top 1,500 differential proteins by the hurdle model as their place in the ranking is taken by identifiers with a strong DD. A similar analysis for the 500 and 1,000 most differential proteins for each method is given in Supplementary Fig. 13.

**Figure 2.**
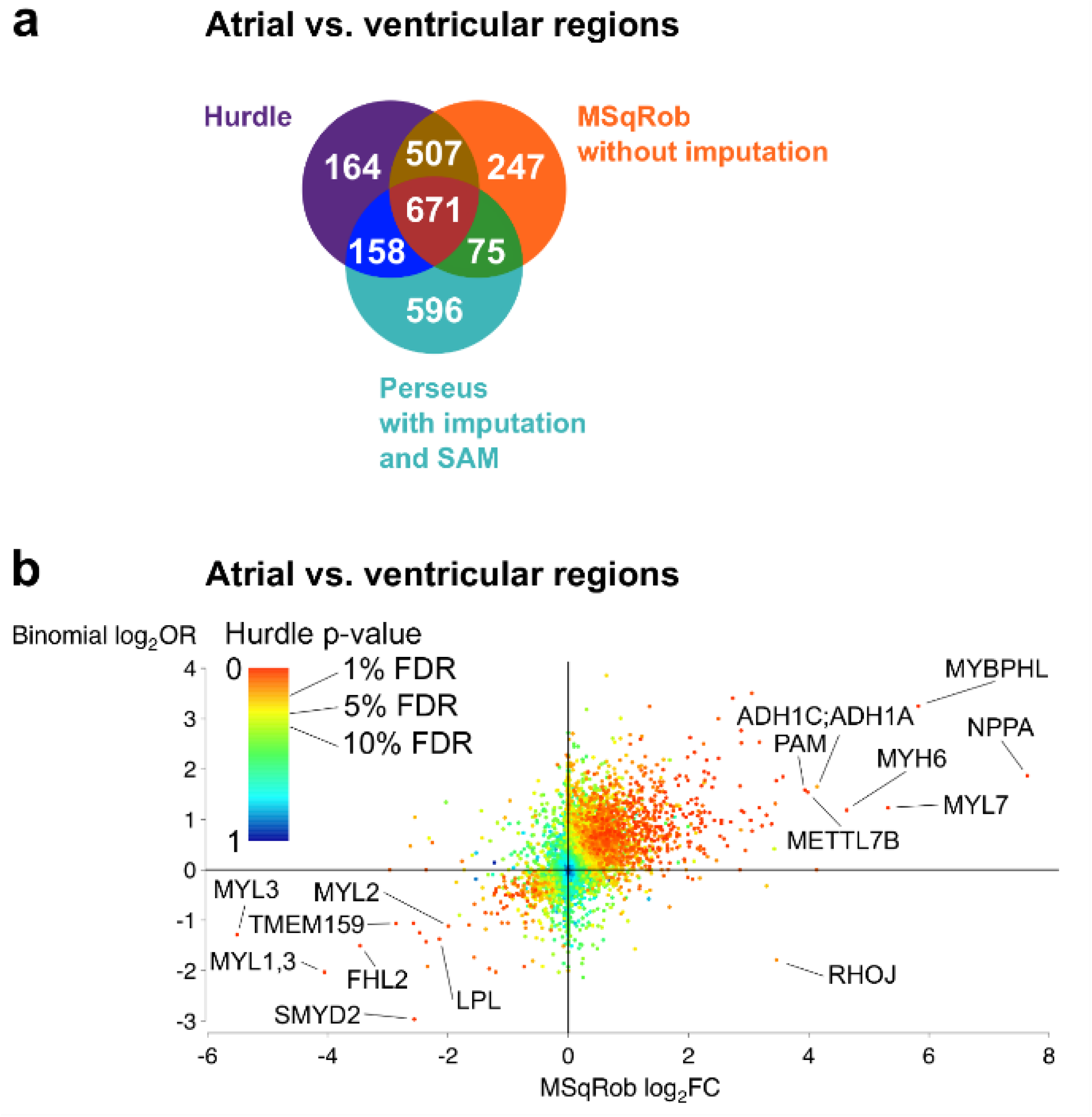
The hurdle model detects both DD and DA proteins. (a) The Venn diagram shows the overlaps in the 1,500 most significant proteins when contrasting atrial (average of left atrium, LA; right atrium, RA; atrial septum, SepA) to ventricular regions (average of left ventricle, LV; right ventricle, RV; ventricular septum, SepV) in the HEART dataset between hurdle, MSqRob without prior imputation, and Perseus with imputation and SAM as implemented by Doll *et al.* (b) Proteins that are significant with the hurdle model often display a strong positive association between FC and OR. Here, log_2_ FCs are plotted as a function of log_2_ ORs in the atrial vs. ventricular comparison for all 6,489 proteins with an FC estimate. Proteins are colored by p-value. Common FDR cut-offs are indicated on the color bar.

For our novel approach we also observe a strong association between the OR and FC estimates (Fig. 2 b). Out of the 1,765 hurdle identifiers that are significant at the 5% FDR level, 397 were significantly DD, 815 significantly DA, 440 both significantly DD and DA, and 113 could not be attributed to either DA or DD in our *post hoc* analysis. All 440 identifiers declared both DD and DA have a FC and OR estimate in the same direction (i.e. both up or both down).

As in the original publication, comparing the left to the right ventricle did not render any significant proteins. However, when comparing the left to the right atrium, we found eight significant proteins compared to only three in the original publication (Supplementary Table 3). The higher abundance of the muscle contraction protein myotilin^22^ in the left atrium was confirmed by both methods, but the hurdle model additionally indicates the heavy myosin chain MYH7 as DA. Moreover, the hurdle model also flags the voltage-dependent calcium channel CACNA2D2 and BMP10, an essential factor for embryonal cardiac development^23^. Finally, SERINC3 and PNMA1, which were declared DA by Doll *et al.* actually come with very limited evidence for DA and their significance is likely driven by the inherent randomness of PI (Supplementary Table 4).

## CONCLUSIONS

In summary, we demonstrated the negative impact of imputation in protein quantification, and the potential flaws it induces in the downstream analysis when imputation assumptions are violated. We therefore propose a hurdle approach that can handle missingness without having to resort to imputation and its associated assumptions by leveraging both an intensity-based approach (MSqRob) and a count-based quasi-binomial approach. Our hurdle approach outcompetes state-of-the-art methods with and without imputation in the presence of both strong missingness as well as when missingness is limited. Moreover, our hurdle method continues to provide valid inference when peptides are absent in one condition. It is therefore a highly promising approach to detect the sudden appearance of post-translationally modified peptides in protein regulation alongside more traditional differential protein expression.

## Supporting information

supplementary_informationHurdle.pdf

## ASSOCIATED CONTENT

### Supporting Information

**supplementary_informationHurdle.pdf:** an extended methods section and all supplementary figures. (PDF)

**supplementary_table1.xlsx:** Overview of the projects and files used in Fig. 1a together with the number of missing values, the number of observed values and the percentage of missing values in each file. (XLSX)

**supplementary_table2.xlsx:** Overview of the relevant protein-level output from the hurdle model (sheet 2), the Pereus with imputation and SAM approach used by Doll *et al.* (sheet 3), MSqRob (sheet 4) and the quasibinomial model (sheet 5) when comparing the atrial to the ventricular regions in the HEART dataset. (XLSX)

**supplementary_table3.xlsx:** Overview of the log_2_ FC, log_2_ OR, χ^2^ statistics, p-values and Benjamini-Hochberg-corrected p-values (q-values) when comparing the left to the right atrium with the hurdle model. (XLSX)

**supplementary_table4.xlsx:** Overview of the data underlying proteins SERINC3, PNMA1, ACSM2A, PDE7B and PNPLA7. (XLSX)

**supplementary_table5.xlsx:** Overview of the mock treatment levels and the condition-lab combinations for each MS run in the mock analysis. (XLSX)

## Author Contributions

L.G. contributed R code, analyzed the data and wrote the paper, A.S. contributed R code and wrote the paper, K.G. and L.M. wrote the paper. L.C. contributed R code, conceived the idea and wrote the paper.

## Notes

The authors declare no competing interests.

## ACKNOWLEDGEMENTS

L.G. is supported by a Ph.D. grant from the Flanders Innovation & Entrepreneurship agency, Flanders (Agentschap Innoveren & Ondernemen – Vlaanderen) entitled ‘Differential proteomics at peptide, protein and module level’ (141573). L.M. acknowledges funding from the Research Foundation Flanders (FWO) under Grant number G042518N. We also thank the students of the Statistical Genomics course, 2016/2017, Ghent University, who assisted us in assessing an initial implementation of the quasi-binomial regression component for their project work.

---

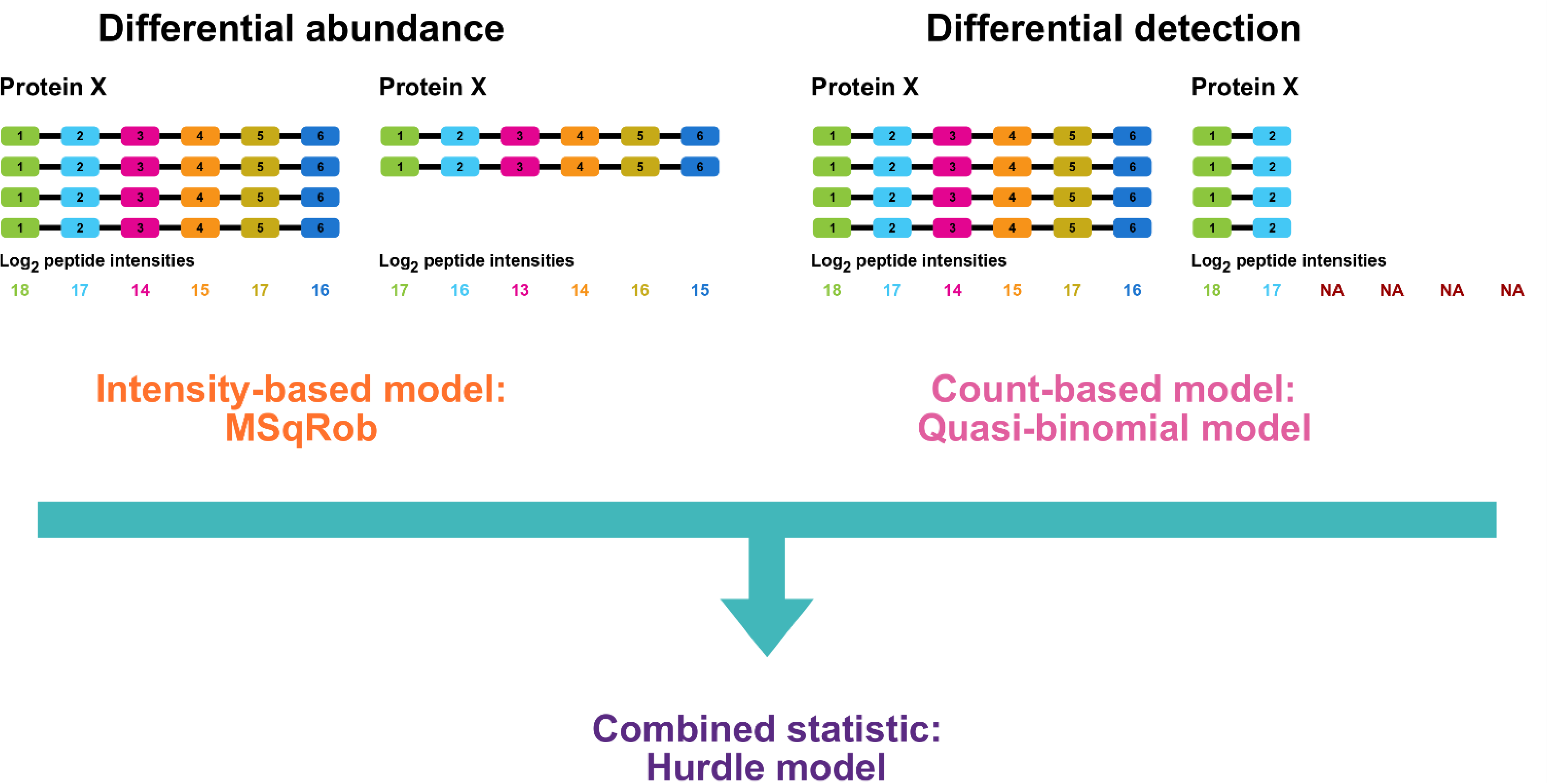

