## supplementary_informationHurdle.pdf for "MSqRob takes the missing hurdle: uniting intensity- and count-based proteomics"

### Supplementary Information

#### Extended methods

For the CPTAC study 6[1], we made use of spike-in conditions A (0.25 fmol UPS1 proteins/µL), B (0.74 fmol UPS1 proteins/µL) and C (2.22 fmol UPS1 proteins/µL), each analyzed on an LTQ-Orbitrap at site 86, an LTQ-Orbitrap O at site 65 and an LTQ-Orbitrap W at site 56. All samples were run in technical triplicate, leading to a total of 27 samples.

The data were searched with MaxQuant 1.6.1.0 as described in the online methods. All settings were kept at their defaults, except for the following:

- MaxQuant was set to perform matching between runs with a match time window of 0.7 min and an alignment time window of 20 min. We also enabled the matching of unidentified features.

- For protein quantification, we set MaxQuant to perform LFQ with a minimal ratio count of 2, a minimum of 3 neighbors and 6 neighbors on average. FastLFQ was enabled.

- As the proteins were alkylated prior to digestion, “carbamidomethyl (C)” is set as a fixed modification. “Acetyl (Protein N-term)”, “Oxidation (M)” and “Gln-pyro-Glu” were set as random modifications.

- We used both the fixed modification and the three random modifications for protein quantification, as there is no differential modification in the CPTAC study, the only difference between samples is the amount of UPS1 proteins.

Scripts to reproduce the plots and tables given in this document are available from: https://github.com/statOmics/MSqRobHurdlePaper.

#### Imputation can greatly distort the distribution of the observed values

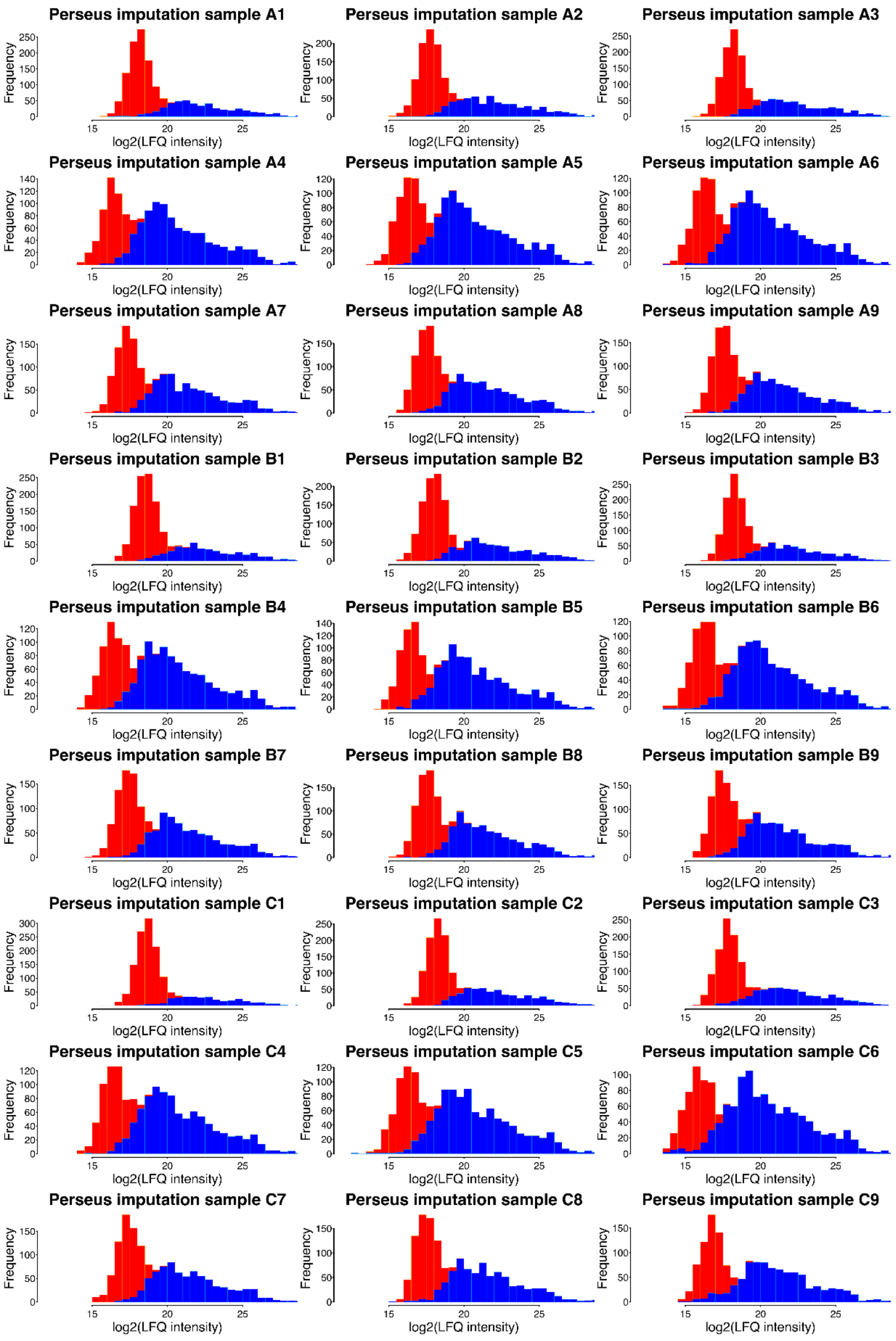


**Supplementary Figure 1.** Perseus imputation breaks down in the presence of many missing values. The histograms show the frequencies of the **log$_2$-transformed LFQ protein intensities** for each sample of the CPTAC dataset after imputing with Perseus. The distributions of the imputed values (red) differ from what would reasonably be expected based on the distribution of the observed values (blue).

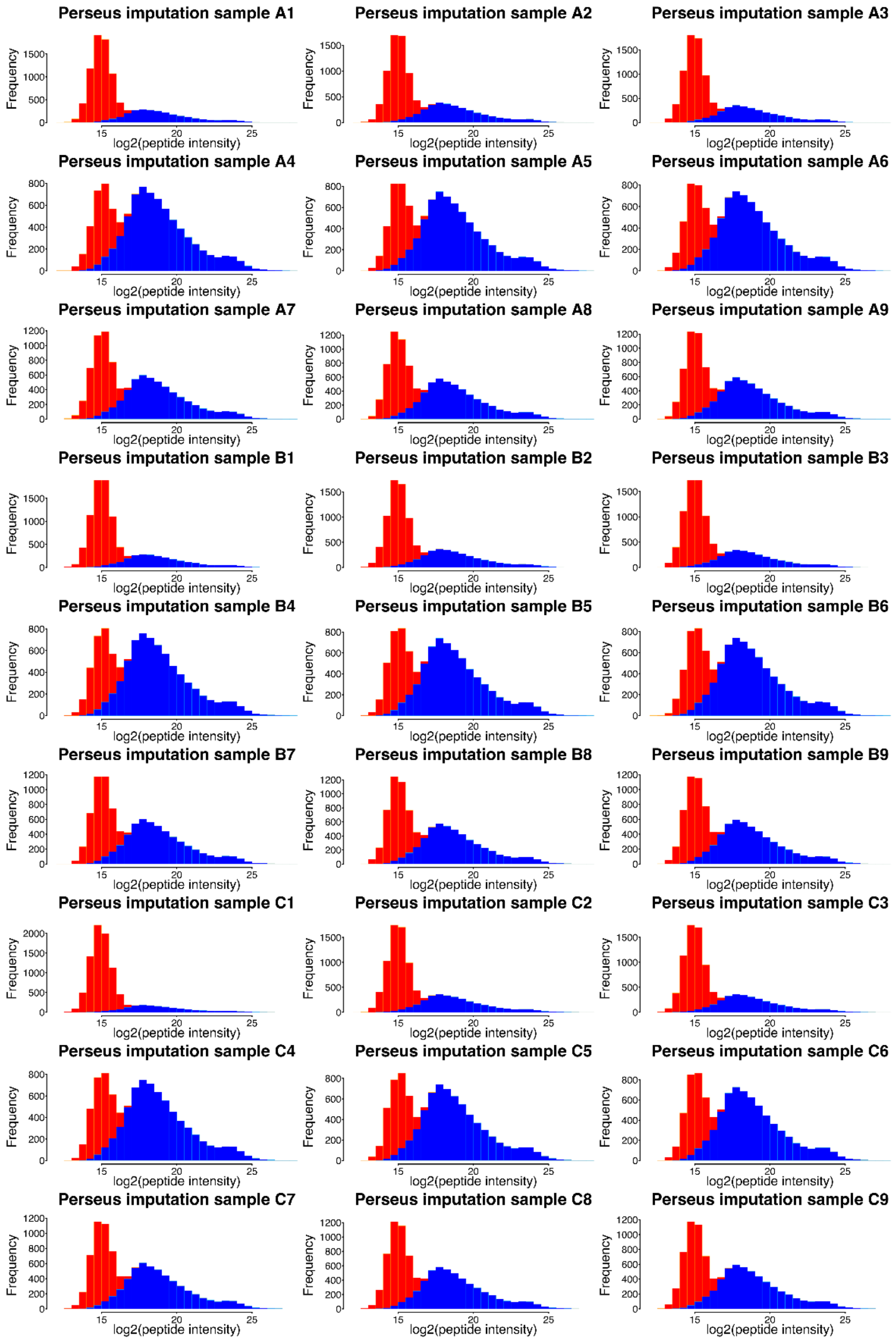


**Supplementary Figure 2.** Perseus imputation breaks down in the presence of many missing values. The histograms show the frequencies of the **$\log_2$-transformed peptide intensities** for each sample of the CPTAC dataset after imputing with Perseus. The distributions of the imputed values (red) differ from what would reasonably be expected based on the distribution of the observed values (blue).

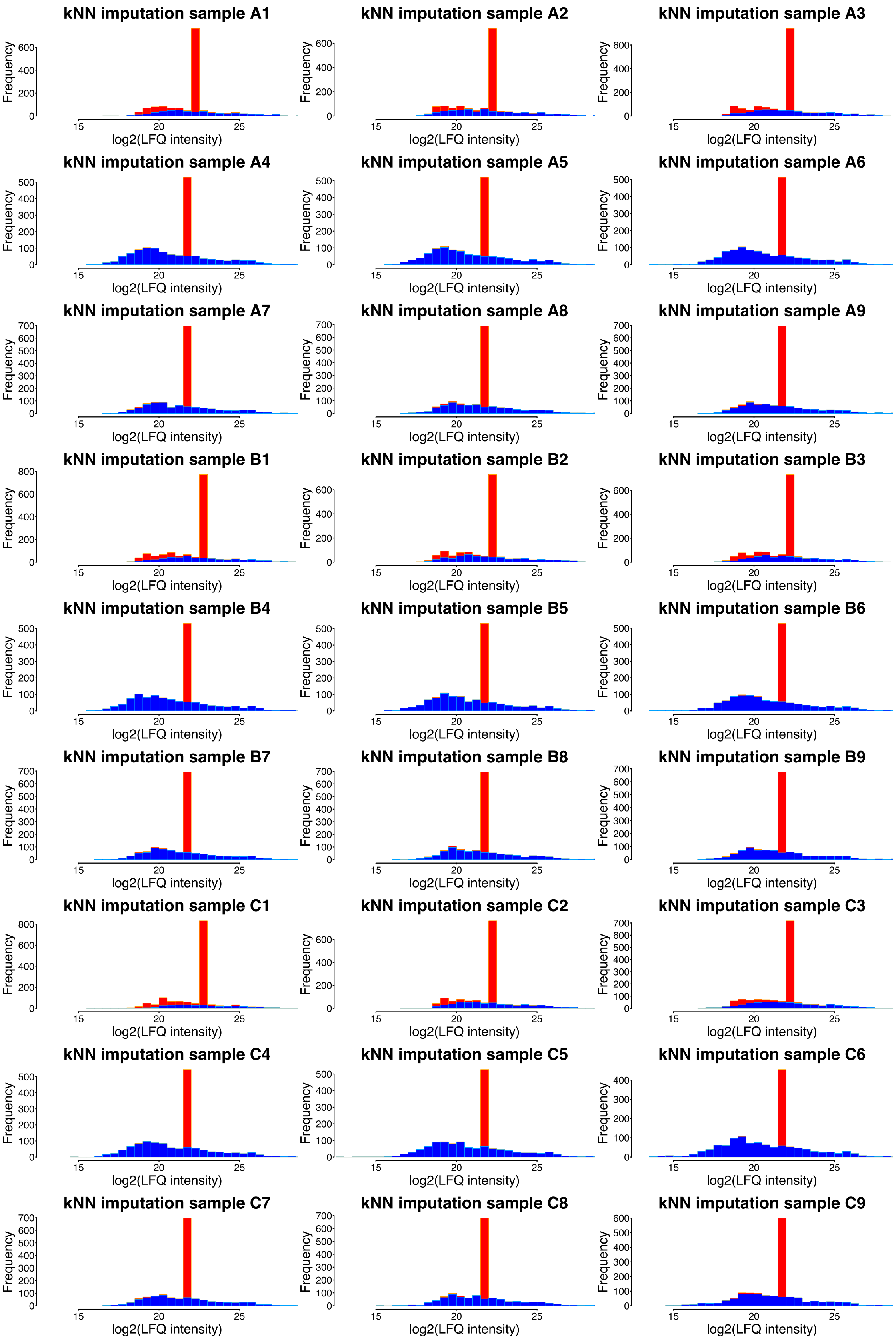


**Supplementary Figure 3.** k-nearest neighbors (kNN) imputation breaks down in the presence of many missing values. The histograms show the frequencies of the **$\log_2$-transformed LFQ protein intensities** for each sample of the CPTAC dataset after imputing with kNN ($k$ = 10). The distributions of the imputed values (red) differ from what would reasonably be expected based on the distribution of the observed values (blue).

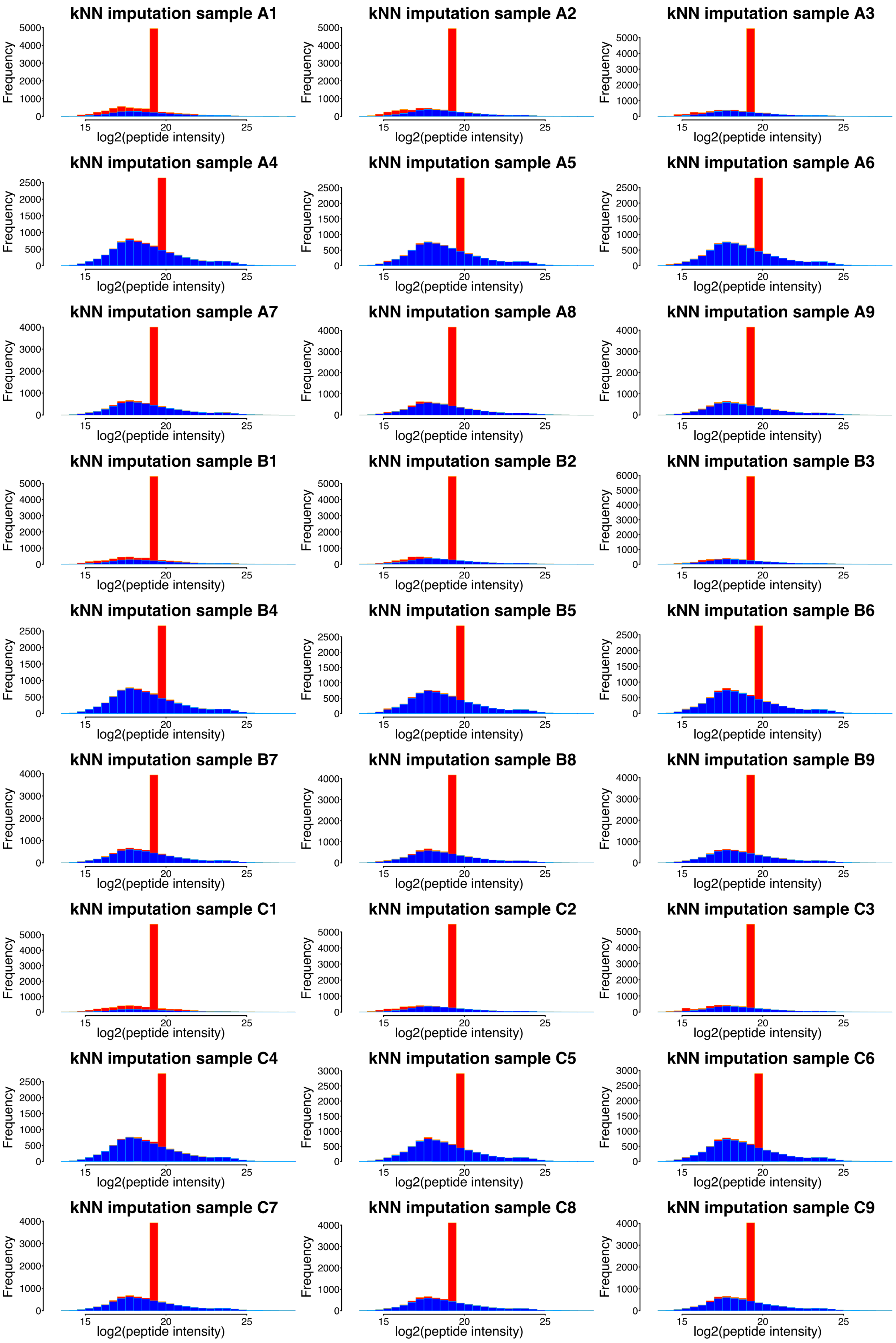


**Supplementary Figure 4.** k-nearest neighbors (kNN) imputation breaks down in the presence of many missing values. The histograms show the frequencies of the **$\log_2$-transformed peptide intensities** for each sample of the CPTAC dataset after imputing with kNN ($k = 10$). The distributions of the imputed values (red) differ from what would reasonably be expected based on the distribution of the observed values (blue).

#### The hurdle model outperforms other methods

Below, we show figures similar to Fig. 1c and 1d, but with addition of the performance of "MSstats with Inf values", "MSqRob with kNN imputation" and "ProPCA with limma". "MSstats with Inf values" is the MSstats approach where proteins for which an FC could not be determined due to an insufficient number of peptides in one condition are assigned an infinite fold change and an FDR of 0. "MSstats without Inf values" is the standard MSstats approach where proteins without an FC estimate are filtered out. "ProPCA with limma" is the combination of ProPCA to summarize intensities and spectral counts into one value[2] with limma for statistical inference[3,4]. We could not reproduce IDPQuantify's performance because that method has only been described, but not implemented[5].

MSqRob with kNN imputation performs very poorly for every comparison of the CPTAC dataset. MSqRob with PI always outperforms MSstats. MSqRob without imputation outperforms MSstats in comparisons B vs. A and C vs. B. Only in comparison C (highest spike-in concentration) vs. A (lowest spike-in concentration), MSstats without Inf values narrowly outperforms MSqRob without imputation. This is very likely due to the fact that the default MSstats pipeline uses an imputation method with a "missingness by low abundance" assumption, an assumption that is valid for the UPS1 proteins in that comparison. MSstats with Inf values performs poorly compared to MSstats without Inf values. Indeed, the naive approach where every protein with missing values is declared significant generates too many false positive hits, as low-abundant proteins can easily go missing in one condition due to random chance.

As the difference in peptide counts for UPS1 proteins is expected to be the largest in C vs. A, the quasibinomial model experiences a remarkable performance boost. In this comparison, MSqRob with PI outperforms MSqRob without imputation. Similarly, there is a boost in performance in Perseus with imputation and MSstats without Inf values in this condition. This is because Perseus' and MSstats' underlying assumptions of missingness due to low abundance are now correct for the UPS1 proteins: there will be relatively few missing values in the high spike-in condition C that will be imputed with low-intensity values.

Also note that the false discovery rate (FDR) is not controlled for most methods in C vs. A. This is very likely due to the effects of ionization competition in the CPTAC dataset that become apparent when comparing very high to very low spike-in conditions [4,6].

In comparison C vs. B, UPS1 spike-in concentrations are relatively high in both conditions. Therefore, a larger fraction of the missing values in these UPS1 proteins is independent of the true protein abundance. Thus, as most UPS1 peptides will be identified in both conditions, the quasibinomial model, which assesses differential detection probabilities, performs poorly. However, despite this poor performance of its quasibinomial component, the performance of the hurdle model remains on par with MSqRob without imputation.

The hurdle model outperforms all other methods in all three comparisons without the need for any imputation.

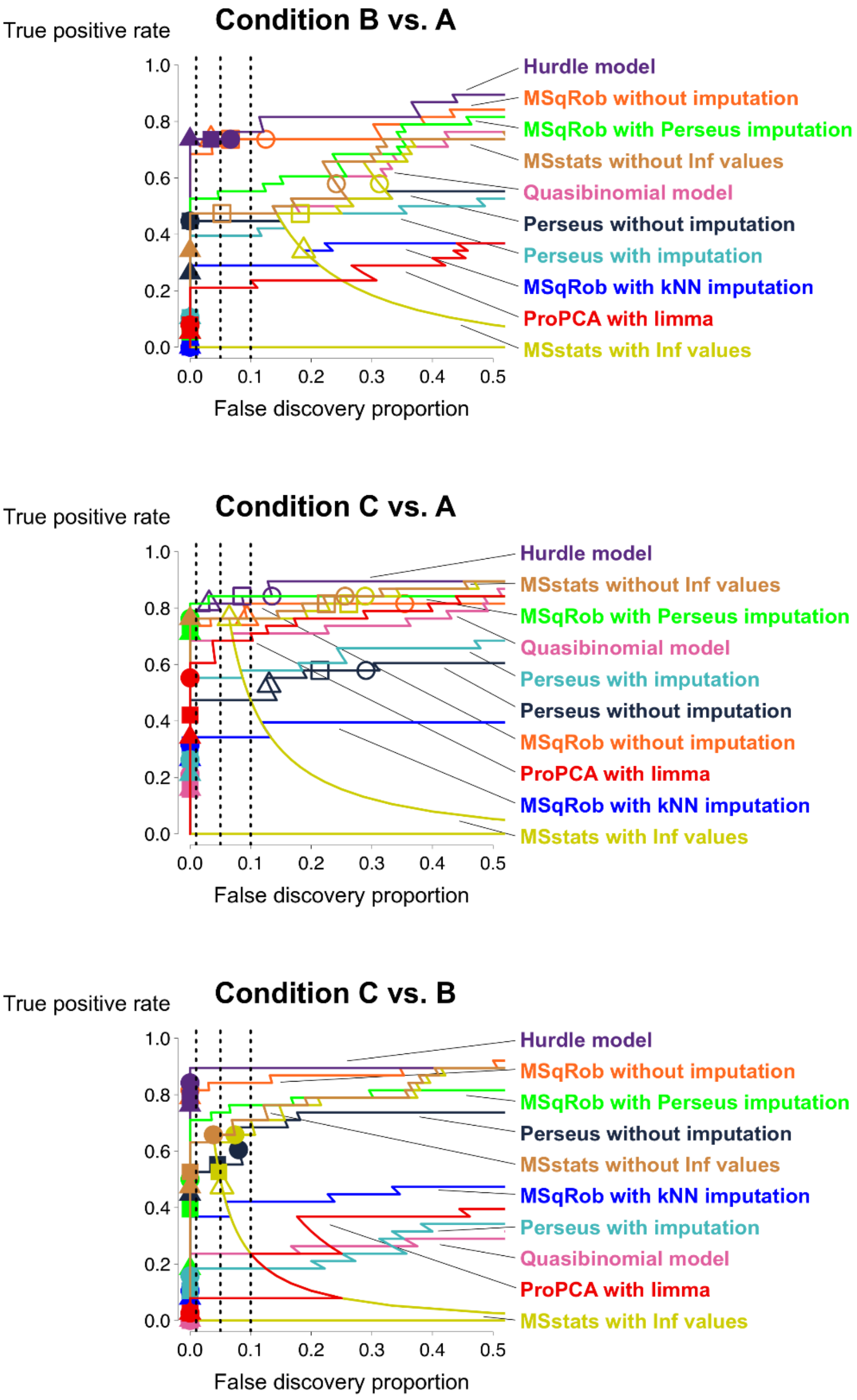


**Supplementary Figure 5.** For the comparisons B vs. A (top), C vs. A (middle) and C vs. B (bottom) in CPTAC dataset, the true positive rate (i.e. the fraction of true positive UPS1 proteins flagged as DA) is plotted as a function of the false discovery proportion (i.e. the fraction of false positive yeast proteins in the total number of DA proteins). Triangles, squares and circles denote the estimated FDR cut offs at 1%, 5% and 10% respectively. Symbols are closed when the FDR is controlled at the given level and open when the FDR is not controlled. “MSstats with Inf values” denotes a default MSstats 3.12.2 pipeline where proteins with an infinite fold change are seen as the most significant hits (FDR set to 0). “MSstats without Inf values” is the same pipeline where proteins with an infinite fold were removed from the data. Here, we used our own preprocessing pipeline as described in the methods section as a reference: any proteins found with MSstats or Perseus that were not present after our preprocessing were removed from the data.

#### The hurdle model outperforms other methods, independent of preprocessing

The two UPS1 proteins, P01112ups and P09211ups, which were removed during our preprocessing, were significant in some MSstats comparisons, but not in Perseus (all p-values = 1). Hence, we decided to redo the analysis with proteins found after preprocessing with MSstats as a basis, to rule out any preprocessing-related effects.

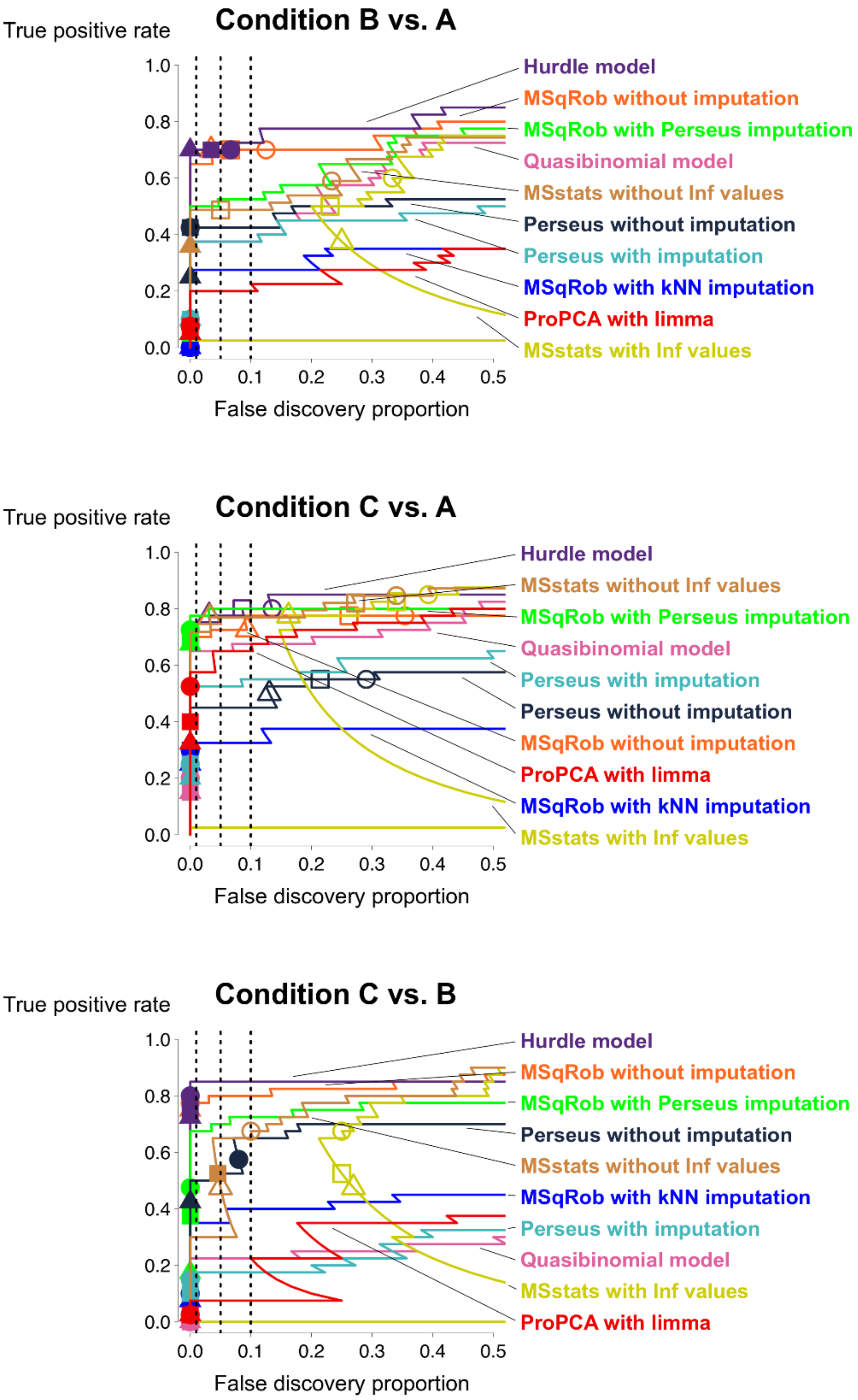


**Supplementary Figure 6.** For the comparisons B vs. A (top), C vs. A (middle) and C vs. B (bottom) in CPTAC dataset, the true positive rate (i.e. the fraction of true positive UPS1 proteins flagged as DA) is plotted as a function of the false discovery proportion (i.e. the fraction of false positive yeast proteins in the total number of DA proteins). Triangles, squares and circles denote the estimated FDR cut offs at 1%, 5% and 10% respectively. Symbols are closed when the FDR is controlled at the given level and open when the FDR is not controlled. “MSstats with Inf values” denotes a default MSstats 3.12.2 pipeline where proteins with an infinite fold change are seen as the most significant hits (FDR set to 0). “MSstats without Inf values” is the same pipeline where proteins with an infinite fold were removed from the data. Here, we used the MSstats preprocessing pipeline as a reference: any proteins found with hurdle, MSqRob or Perseus that were not present after MSstats’ preprocessing were removed from the data.

#### Intensities and counts provide complementary information for detecting DA

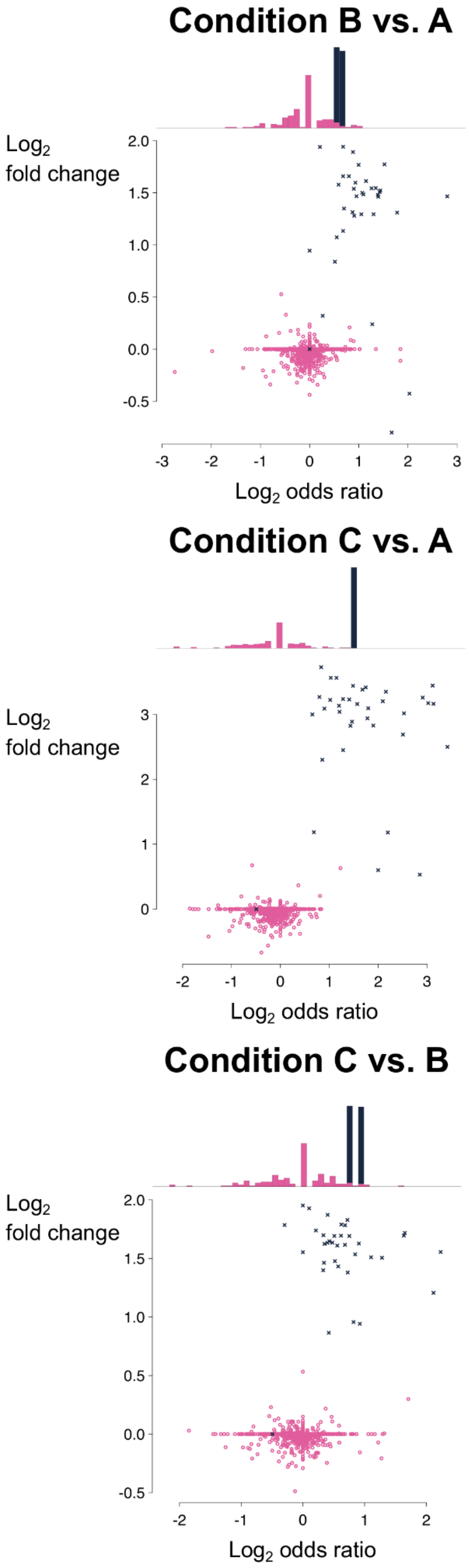


**Supplementary Figure 7.** Intensities and counts provide complementary information for detecting differentially abundant proteins. The scatter plot shows MSqRob's intensity-based estimated $\log_2$ fold change on the vertical axis and the quasibinomial model's count-based estimated $\log_2$ odds ratio on the horizontal axis for all proteins in the CPTAC dataset for comparisons B vs. A, C vs. A and C vs. B for which a $\log_2$ fold change could be estimated. Histogram: marginal distribution for the $\log_2$ odds ratio of the 158 (C vs. A) or 159 (B vs. A and C vs. B) yeast proteins and 2 UPS1 proteins (all comparisons) for which no $\log_2$ fold change could be estimated. UPS1 proteins (TP) are dark blue crosses, while yeast proteins (FP) are pink circles.

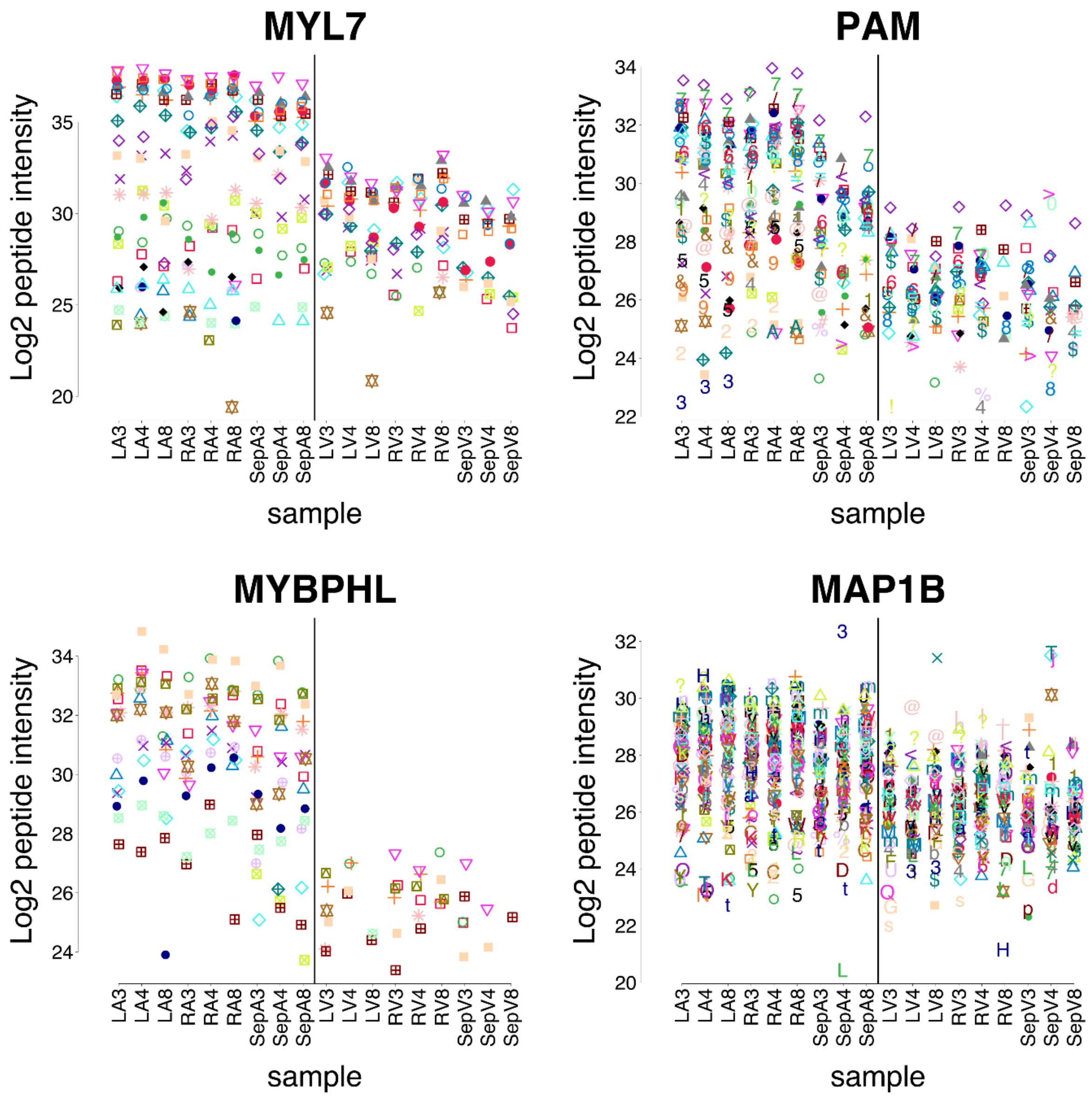


**Supplementary Figure 8.** In the subset of the 1500 most significant gene identifiers for the atrial to the ventricular proteome comparison in the HEART dataset, proteins that are identified by Perseus with imputation and SAM, MSqRob and the hurdle model often have large fold changes and many identified peptides, as exemplified by proteins MYL7, PAM, MYBPHL and MAP1B. The plots show the preprocessed $\log_2$-peptide intensities per sample. Each different combination of color and shape denotes a different peptide sequence. LA: left atrium, RA: right atrium, SepA: atrial septum, LV: left ventricle, RV: right ventricle, SepV: ventricular septum. 3, 4 and 8 indicate the three different healthy hearts used in the study.  Similar plots for all 671 proteins identified by Perseus with imputation and SAM, MSqRob, and the hurdle model in the first 1,500 gene identifiers of the HEART dataset can be found at: https://github.com/statOmics/MSqRobHurdlePaper/raw/master/sign_protein_plots_PXD006675/overlap_all.pdf.

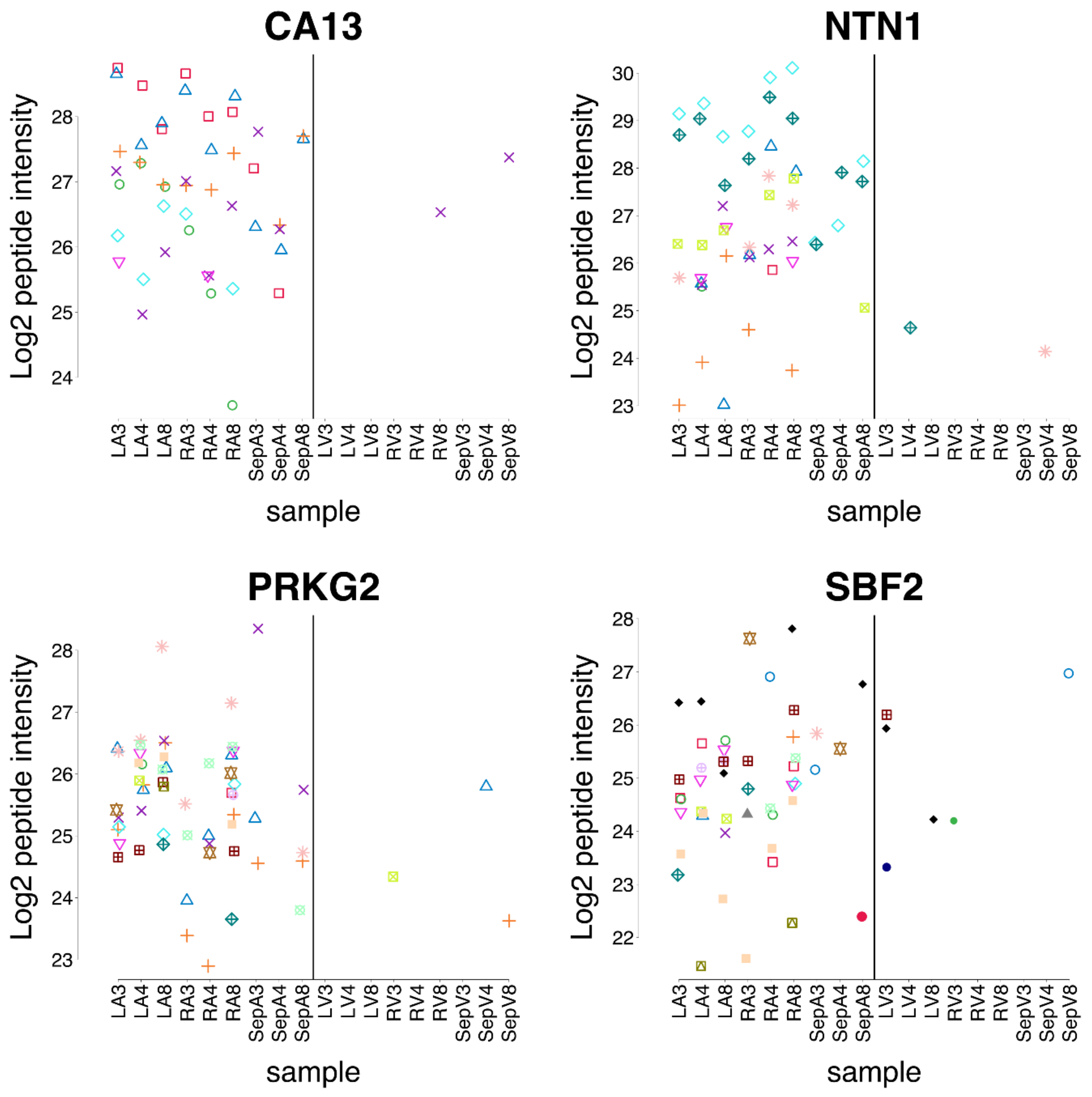


**Supplementary Figure 9.** In the subset of the 1500 most significant gene identifiers for the atrial to the ventricular proteome comparison in the HEART dataset, proteins that are identified by Perseus with imputation and SAM, and the hurdle model, but not MSqRob, often show a strong differential detection with little peptides identified in one of both heart chambers, as exemplified by proteins CA13, NTN1, PRKG2 and SBF2. The plots show the preprocessed $log_2$-peptide intensities per sample. Each different combination of color and shape denotes a different peptide sequence. LA: left atrium, RA: right atrium, SepA: atrial septum, LV: left ventricle, RV: right ventricle, SepV: ventricular septum. 3, 4 and 8 indicate the three different healthy hearts used in the study. Similar plots for all 158 proteins identified by Perseus with imputation and SAM, and the hurdle model, but not MSqRob, in the first 1,500 gene identifiers of the HEART dataset can be found at: https://github.com/statOmics/MSqRobHurdlePaper/raw/master/sign_protein_plots_PXD006675/overlap_hurdle_Perseus.pdf.

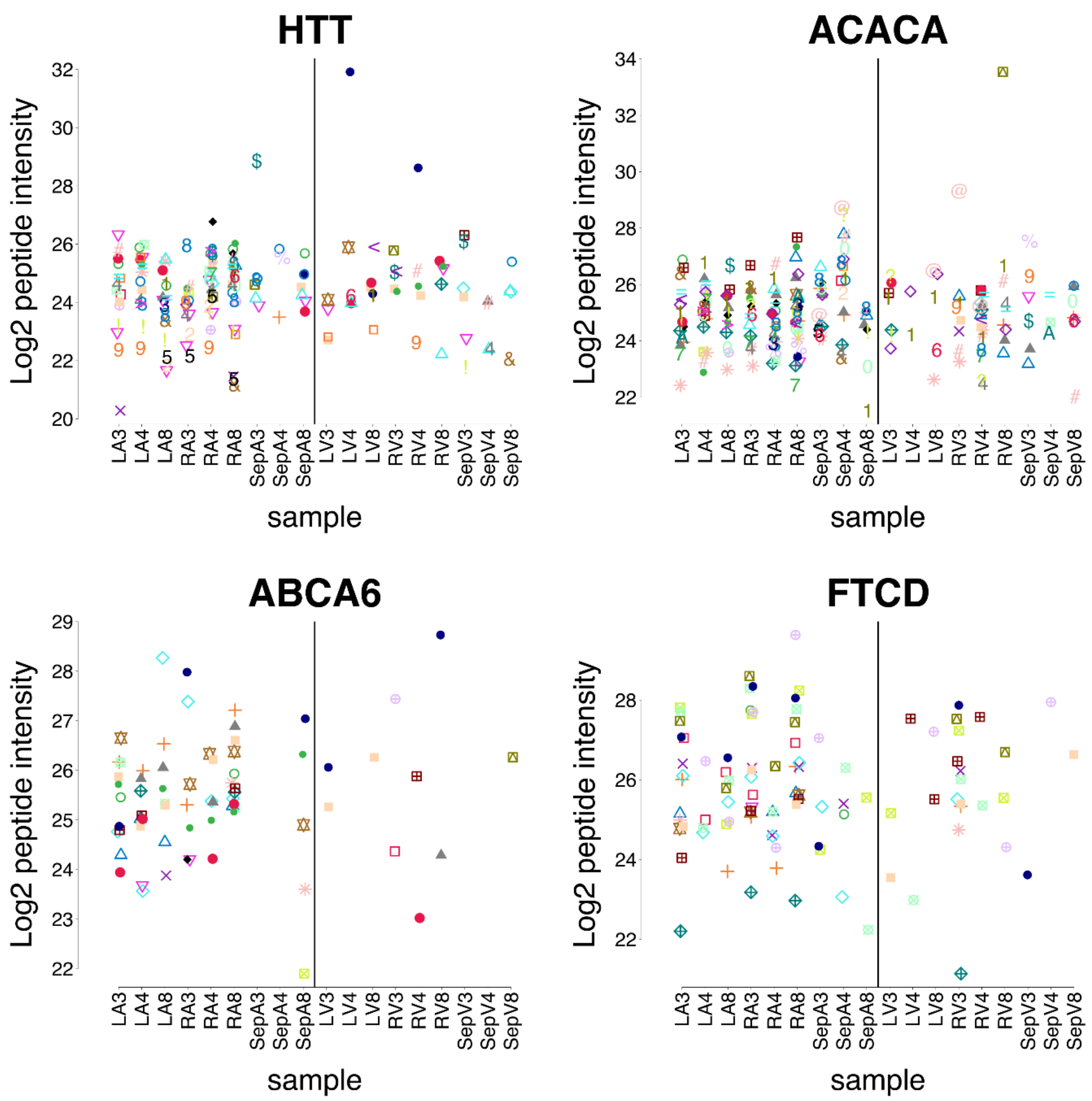


**Supplementary Figure 10.** In the subset of the 1500 most significant gene identifiers for the atrial to the ventricular proteome comparison in the HEART dataset, proteins that are identified by the hurdle model, but not by MSqRob and Perseus with imputation and SAM, often show a strong differential detection, but have few to no samples wherein no peptides were detected, as exemplified by proteins HTT, ACACA, ABCA6 and FTCD. The plots show the preprocessed $log_2$-peptide intensities per sample. Each different combination of color and shape denotes a different peptide sequence. LA: left atrium, RA: right atrium, SepA: atrial septum, LV: left ventricle, RV: right ventricle, SepV: ventricular septum. 3, 4 and 8 indicate the three different healthy hearts used in the study. Similar plots for all 164 proteins identified by the hurdle model, but not by MSqRob and Perseus with imputation and SAM, in the first 1,500 gene identifiers of the HEART dataset can be found at: https://github.com/statOmics/MSqRobHurdlePaper/raw/master/sign_protein_plots_PXD006675/only_hurdle.pdf.

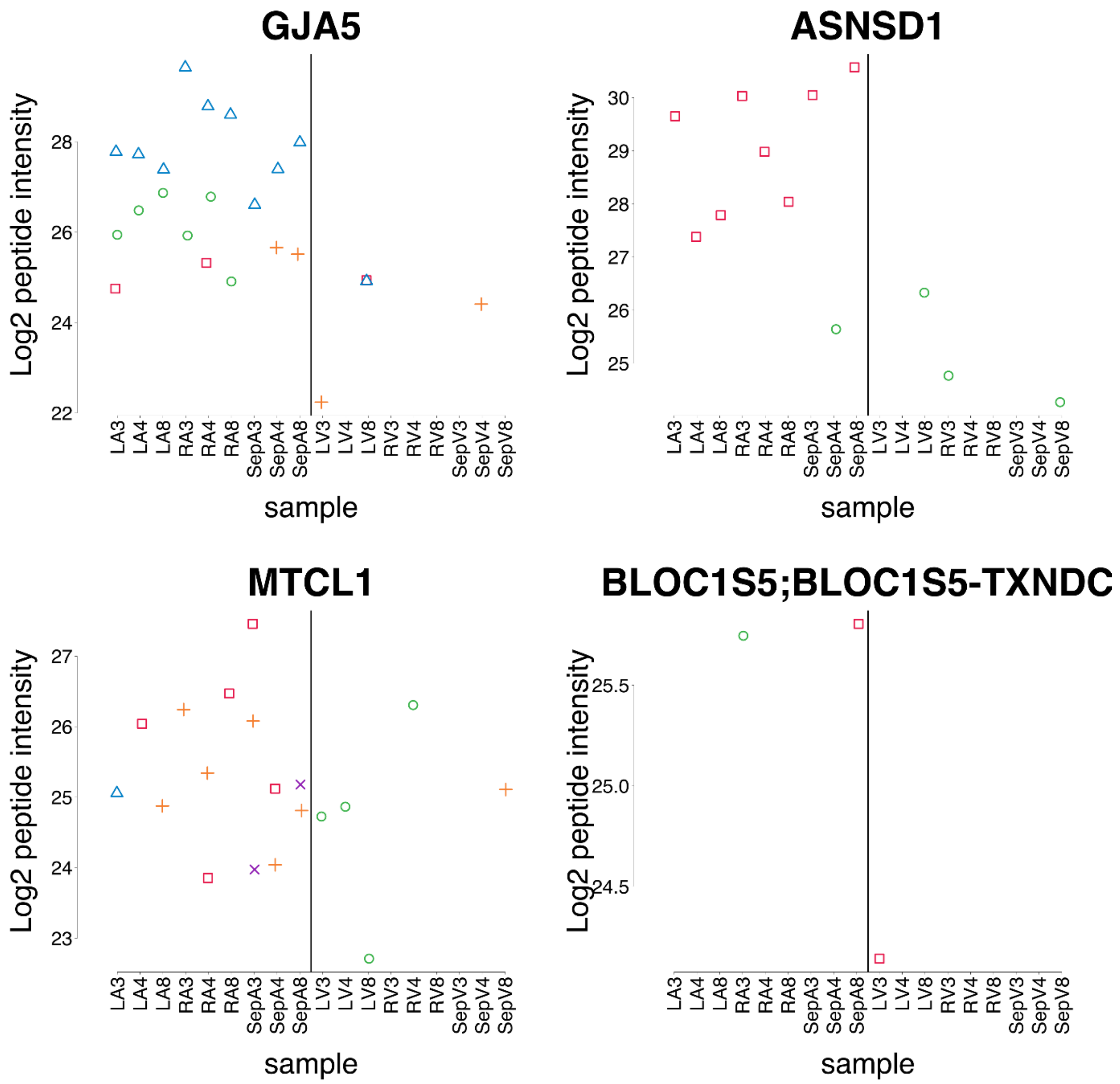


**Supplementary Figure 11.** In the subset of the 1500 most significant gene identifiers for the atrial to the ventricular proteome comparison in the HEART dataset, proteins that are identified by Perseus with imputation and SAM, but not by MSqRob and the hurdle model, often have either rather few peptide identifications, or very small fold changes as exemplified by proteins GJA5, ASNSD1, MTCL1 and BLOC1S5;BLOC1S5-TXNDC5, respectively. The plots show the preprocessed $\log_2$-peptide intensities per sample. Each different combination of color and shape denotes a different peptide sequence. LA: left atrium, RA: right atrium, SepA: atrial septum, LV: left ventricle, RV: right ventricle, SepV: ventricular septum. 3, 4 and 8 indicate the three different healthy hearts used in the study. Similar plots for all 596 proteins identified Perseus with imputation and SAM, but not by MSqRob and the hurdle model, in the first 1,500 gene identifiers of the HEART dataset can be found at: https://github.com/statOmics/MSqRobHurdlePaper/raw/master/sign_protein_plots_PXD006675/only_Perseus.pdf.

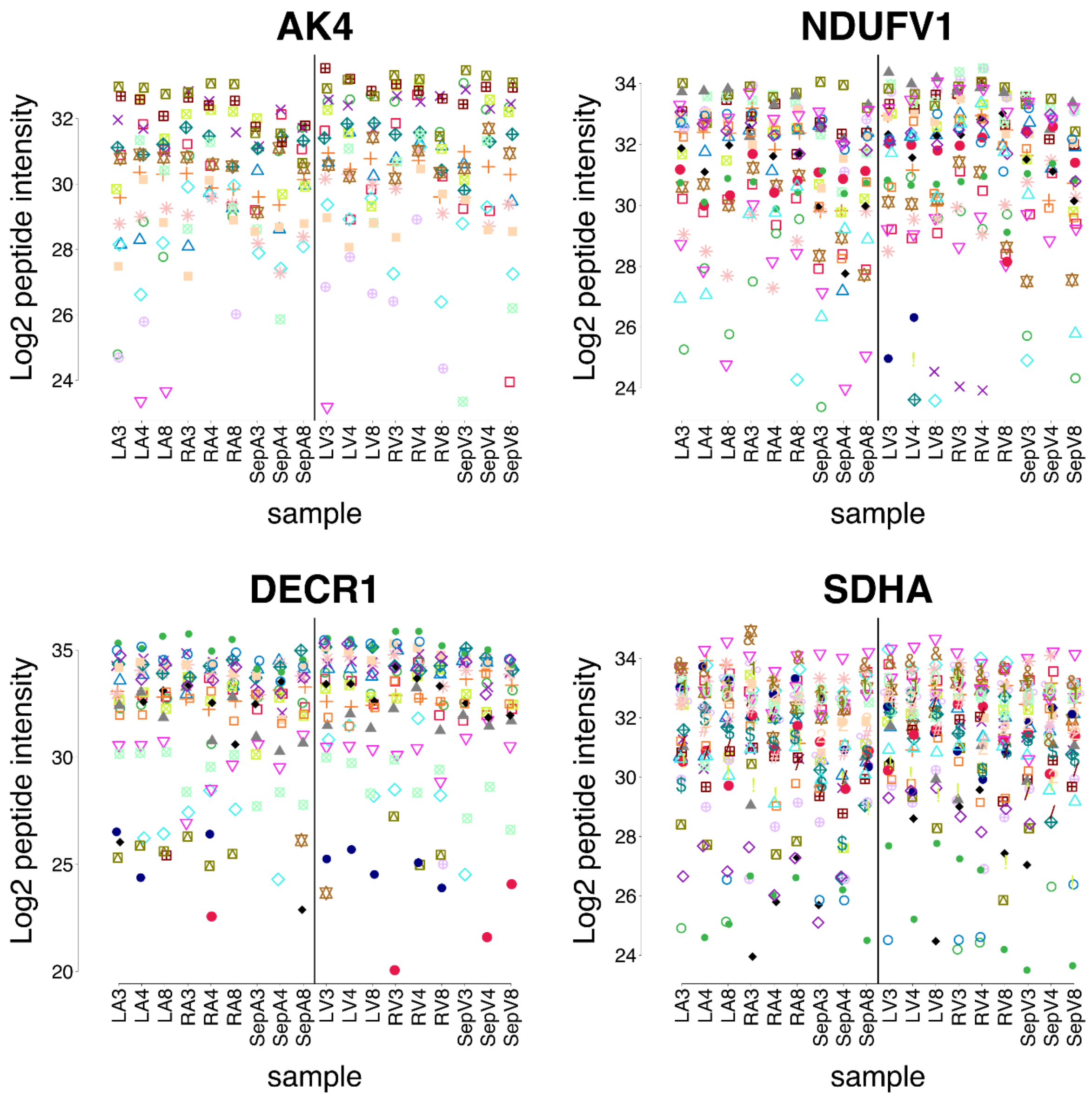


**Supplementary Figure 12.** In the subset of the 1500 most significant gene identifiers for the atrial to the ventricular proteome comparison in the HEART dataset, proteins that are identified by MSqRob, and Perseus with imputation and SAM, but not by the hurdle model often show many identified peptides, but rather small fold changes, as exemplified by proteins AK4, NDUFV1, DECR1 and SDHA. The plots show the preprocessed $\log_2$-peptide intensities per sample. Each different combination of color and shape denotes a different peptide sequence. LA: left atrium, RA: right atrium, SepA: atrial septum, LV: left ventricle, RV: right ventricle, SepV: ventricular septum. 3, 4 and 8 indicate the three different healthy hearts used in the study. Similar plots for all 75 proteins identified by MSqRob, and Perseus with imputation and SAM, but not the hurdle model, in the first 1,500 gene identifiers of the HEART dataset can be found at:
https://github.com/statOmics/MSqRobHurdlePaper/raw/master/sign_protein_plots_PXD006675/overlap_MSqRob_Perseus.pdf.

#### Overlap between different methods

Different preprocessing approaches and some differences in gene identifiers between the peptides.txt file and the proteinGroups.txt file make that not all proteins found with MSqRob/hurdle could be found with Perseus, and vice versa. To keep a fair comparison, we here only considered those proteins found by both approaches.

We showed the overlap between the 1,500 most significantly regulated proteins between the hurdle model, MSqRob without imputation, and Perseus with imputation and SAM (as implemented by Doll *et al.*) in the main manuscript. Here, we show the same Venn diagrams for the 500 and 1,000 most significantly regulated proteins as well as the overlap between all significant proteins at 5% FDR for each approach. As stated before, proteins that could not be found in both preprocessing approaches were filtered out in advance.

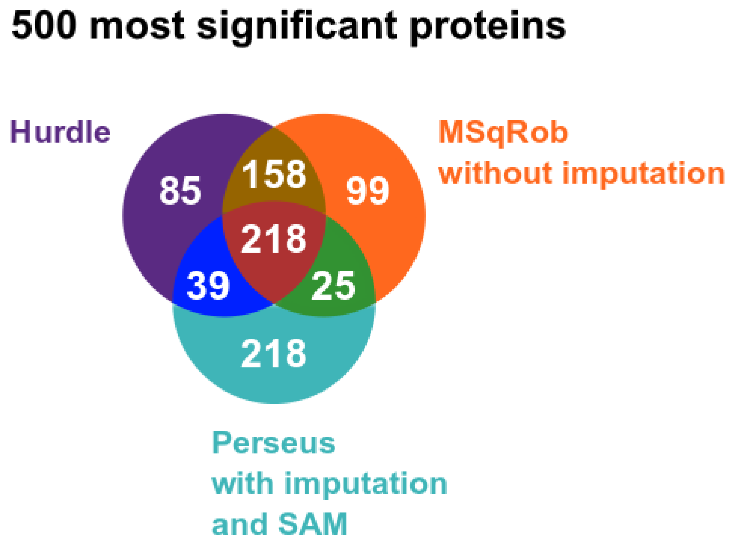


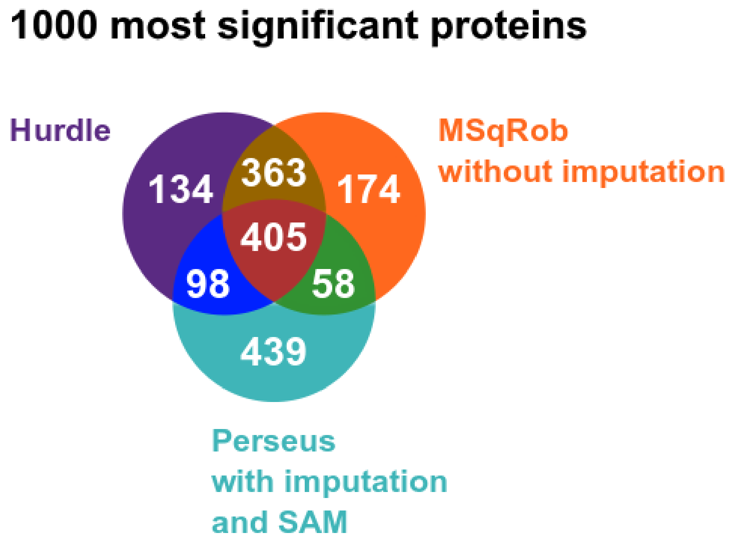


**Supplementary Figure 13.** The Venn diagrams show the overlaps in the 500 most significant proteins (top) and the 100 most significant proteins (bottom) when comparing the atrial to ventricular regions in the HEART dataset between hurdle, MSqRob without prior imputation, and Perseus with imputation and SAM as implemented by Doll *et al.*

#### Comparing the left to the right heart atrium with the original approach of Doll *et al.*

Supplementary Table 4 demonstrates the unpredictable and undesirable effects of Perseus imputation: proteins with very little evidence for DA are declared significant by Doll *et al*., while other proteins in the dataset that have, in our opinion, at least as much (or little) evidence for DA were not declared significant. This is solely due to the inherent randomness of Perseus imputation.

We first show the peptide-level data for SERINC3 and PNMA1, two proteins that were reported to be significantly different at the 5% FDR level in the left to the right heart atrium comparison by Doll *et al*. We then compare this data to the peptide-level data for ACSM2A, PDE7B and PNPLA7, three examples from the dataset with very similar evidence for DA as SERINC3 and PNMA1.

The fact that SERINC3 and PNMA1 were declared DA by Doll *et al*. and those three other proteins were not, again demonstrates the undesirable effects of imputation: depending on the randomly imputed values, a protein can be declared significant or not.

#### On the independence of the z-statistics under the null hypothesis

Under the null hypothesis of no differential abundance, the combined test statistic $\chi^2_{\text{hurdle}}$ $(\text{Eq. } 10)$ will follow a chi-squared distribution with two degrees of freedom if both z-statistics are independent. To assess this assumption, we performed a mock analysis on the repeats of the spike-in conditions within each lab. We also included the combination of spike-in condition and lab as a blocking factor in the analysis (see Supplementary Table 5). This set-up ensures that none of the proteins are differentially abundant.

The results of the three pairwise comparisons, both for the MSqRob and for the quasi-binomial model component of the hurdle model are shown in Fig. 3. and confirm that the estimates for the $\log_2$ FCs and the log ORs are not correlated in the mock analysis.

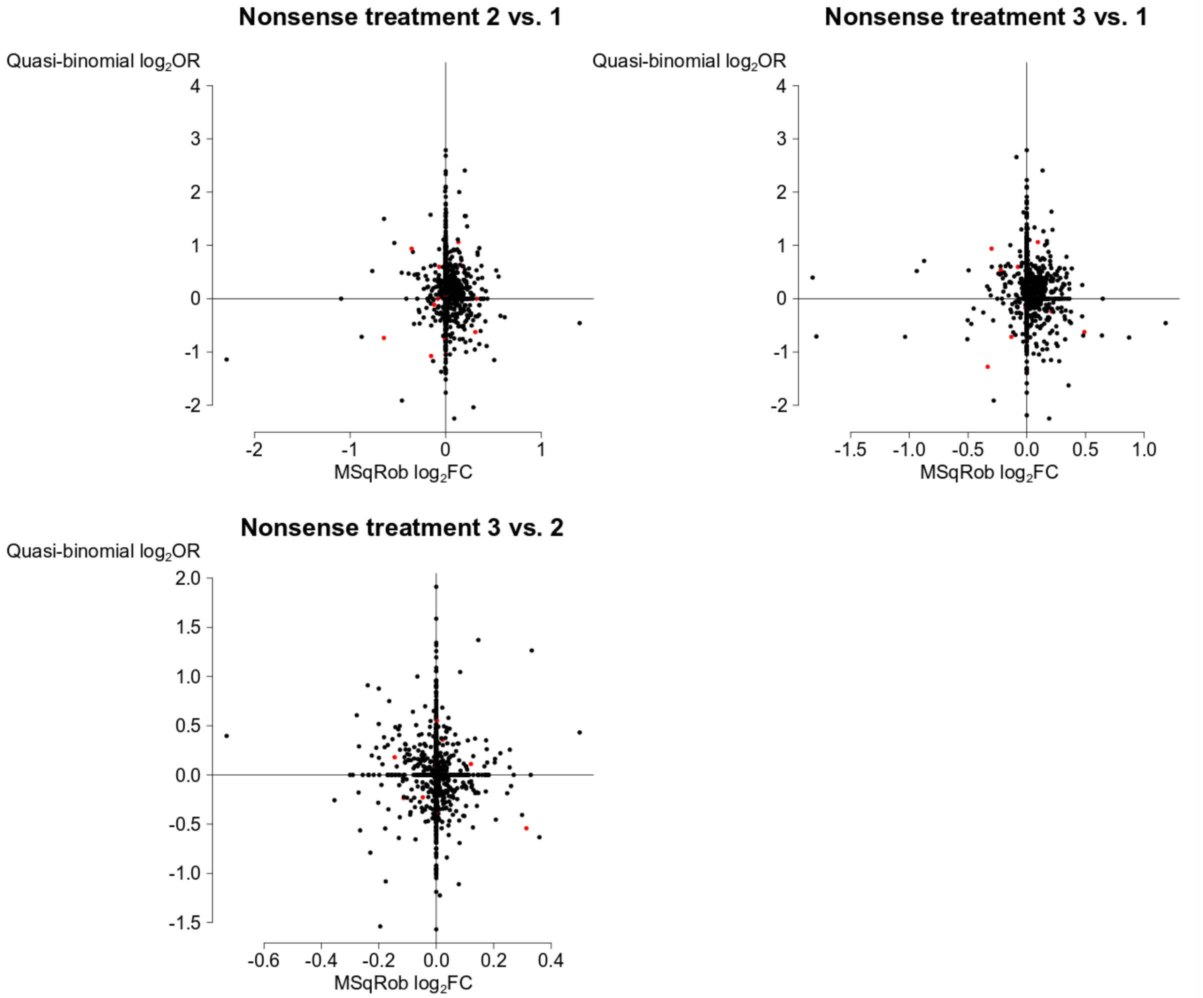


**Supplementary Figure 14.** There seems to be no correlation between the estimates from the MSqRob and quasi-binomial model component. The scatter plots show the $log_2$ FC estimates on the horizontal axis and log OR estimates on the vertical axis of the hurdle model for all proteins in the CPTAC dataset for which a $log_2$ FC could be estimated in all three comparisons of the mock analysis (UPS1 proteins are indicated in red, while yeast proteins are indicated in black).
